# Discovery and characterization of a chemical probe targeting the zinc-finger ubiquitin-binding domain of HDAC6

**DOI:** 10.1101/2023.02.21.525740

**Authors:** Rachel J. Harding, Ivan Franzoni, Mandeep K. Mann, Magdalena M. Szewczyk, Bijan Mirabi, Dominic D.G Owens, Suzanne Ackloo, Alexej Scheremetjew, Kevin A. Juarez-Ornelas, Randy Sanichar, Rachel J. Baker, Christian Dank, Peter J. Brown, Dalia Barsyte-Lovejoy, Vijayaratnam Santhakumar, Matthieu Schapira, Mark Lautens, Cheryl H. Arrowsmith

**Author notes:** Author has updated affiliation: Valence Discovery Inc., 6666 Rue St-Urbain, Suite 200, Montreal, Quebec, H2S 3H1, Canada (Ivan Franzoni). These authors contributed equally (Rachel J. Harding, Ivan Franzoni, Mandeep K. Mann, Magdalena M. Szewczyk, Bijan Mirabi). Deceased July 1, 2021 (Kevin A. Juarez-Ornelas). Corresponding authors – Mark Lautens -, Cheryl H. Arrowsmith.

## Abstract

Histone deacetylase 6 (HDAC6) inhibition is an attractive strategy for treating numerous cancers, and HDAC6 catalytic inhibitors are currently in clinical trials. The HDAC6 zinc-finger ubiquitin-binding domain (UBD) binds free C-terminal diglycine motifs of unanchored ubiquitin polymer chains and protein aggregates, playing an important role in autophagy and aggresome assembly. However, targeting this domain with small molecule antagonists remains an underdeveloped avenue of HDAC6-focused drug discovery. We report SGC-UBD253 (25), a chemical probe potently targeting HDAC6-UBD in vitro with selectivity over nine other UBDs, except for weak USP16 binding. In cells, **25** is an effective antagonist of HDAC6-UBD at 1 µM, with marked proteome-wide selectivity. We identified SGC-UBD253N (**32**), a methylated derivative of **25** which is 300-fold less active, serving as a negative control. Together, **25** and **32** could enable further exploration of the biological function of the HDAC6 UBD and investigation of the therapeutic potential of targeting this domain.

## INTRODUCTION

There are 18 human histone deacetylases (HDACs) which regulate a plethora of important cellular functions by catalyzing the removal of acetyl groups from lysine residues of both histone and non-histone proteins ^1^. HDAC6 is a structurally distinct microtubule-associated cytosolic deacetylase, harboring two tandem catalytic deacetylase domains ^2, 3^ and a zinc-finger ubiquitin-binding domain (UBD) ^4, 5^. The catalytic domains interact with dynein to transport aggregated proteins via microtubules to the aggresome for degradation ^6–8^ whereas the UBD binds free C-terminal diglycine motifs of polyubiquitin or polyISG chains associated with protein aggregates or other cargo ^9, 10^.

Inhibiting HDAC6 function is postulated to have therapeutic benefit in a number of different cancers, neurodegenerative diseases and other pathologies ^11–13^. To date, HDAC6 drug discovery has focused on inhibitors targeting the catalytic activity of this protein, and are currently being tested in the clinic, in some cases in combination with proteasome inhibitors ^14–17^. HDAC6 catalytic inhibitors prevent the deacetylation of microtubules, which disrupts dynein-mediated transport of protein cargos to the aggresome. However, current selective catalytic hydroxamate HDAC6 inhibitors have demonstrated selectivity and toxicity liabilities ^18^. An alternative approach to inhibit or modulate HDAC6 function is to target protein aggregate recognition by the UBD with small molecule antagonists, both in isolation or, in combination with catalytic inhibition.

We previously identified and characterized the first small molecule binders of HDAC6 UBD which can displace the native C-terminal ubiquitin RLRGG peptide from a narrow and deep pocket within this domain ^19^. These compounds have a carboxylate which mimics the C-terminus of the ubiquitin substrate, and an extended aromatic structure which forms π-stacking interactions with W1182 and R1155. Together with our subsequent structure-activity relationship analysis of this chemical series ^20^, we found that it was possible to induce a conformational remodeling of the pocket, to open up an additional side pocket, which we postulated could be exploited to increase potency and selectivity.

We describe the discovery and characterization of **SGC-UBD253** (**25**), a potent antagonist HDAC6-UBD chemical probe, active in cells at 1 µM and selective against other UBDs, albeit with weak activity for USP16. We present the optimization of our previous lead ligand (**1**) ^20^ to improve potency using a fluorescence polarization (FP) C-terminal ubiquitin peptide displacement assay. This led to identification of our probe candidate (**25)**, and a negative control compound (**32**), which is structurally similar to **25** with an additional methyl group which drastically decreases activity. We validated the binding parameters of **25** and **32** with multiple orthogonal biophysical assays and show that the chemical probe (**25)** causes significant activity in cells at 1 µM, whilst the negative control (**32)** is inactive at all concentrations tested. We determined with chemoproteomic approaches that **25** has marked proteome-wide selectivity and that at a working concentration of 3 µM, **25** has significant cellular activity whilst limiting off-target activity for USP16. These tool compounds will enable biological discovery of the functional roles of the UBD of HDAC6 and serve as a foundation for further optimization to develop drugs which target HDAC6-UBD.

## RESULTS AND DISCUSSION

### STRUCTURE−ACTIVITY RELATIONSHIP

We previously reported **1**, a fragment hit, that binds to HDAC6 UBD which we characterized with established FP (K_disp_ ∼ 2.3 μM) and surface plasmon resonance (SPR) assays (K_D_ ∼1.5 μM) ^19, 20^. In the co-crystal structure of **1** with the HDAC6 ubiquitin binding domain (PDB ID: 6CED), the quinazolinone ring is sandwiched between R1155 and W1182 and the carboxylic acid group makes a hydrogen bond between G1154 and Y1184, as well as with R1155. The co-crystal structure also revealed that the N-Methyl group of **1** could be extended into the adjacent pocket to make additional interactions with the residues lining this site (**Figure 1**).

**Figure 1.**
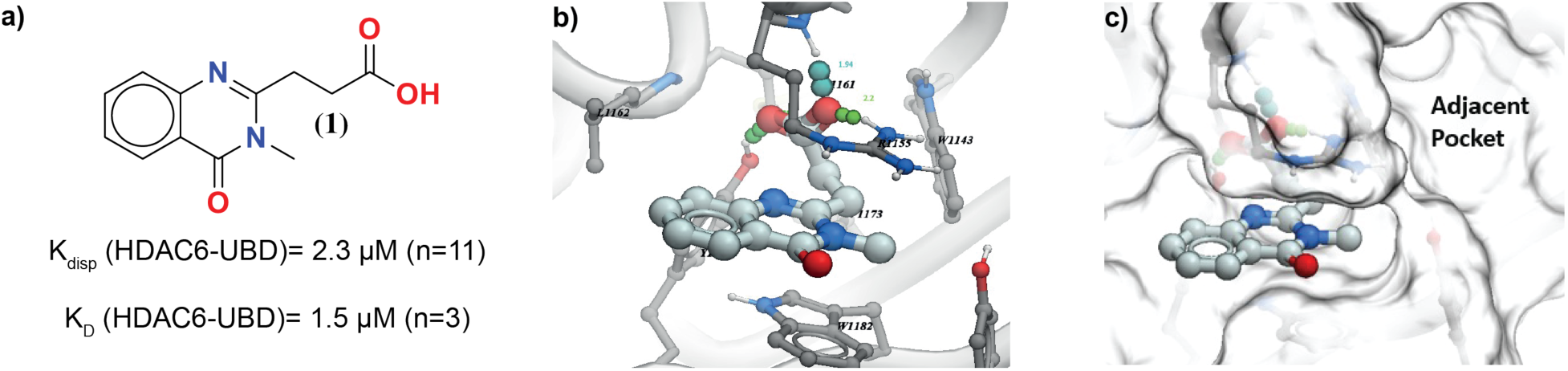
Structure and co-crystal structure of **1** in complex with HDAC6 UBD. a) Structure of compound **1** and associated binding parameters from FP and SPR assays. b) Co-crystal structure of **1** with HDAC6 UBD showing key hydrogen bond interactions, c) Space filled diagram showing adjacent pocket.

To test this hypothesis, we first explored simple *N*-alkyl groups such as cyclopropylmethyl and benzyl groups which could potentially extend into the side pocket, but they led to significant loss of activity (data not shown). Therefore, we explored extended linker groups, starting from the corresponding methyl acetamide derivative (**9**), which was synthesized according to **Scheme 1**. Anthranilic acid (**2a**) was converted into amide (**3a**) which was subsequently treated with succinic anhydride to give **4a**. Cyclization of **4a** in the presence of sodium hydroxide followed by esterification with ethanol gave the ethyl ester protected intermediate (**6a**). Alkylation of **6a** with *N*-methylbromoacetamide (**7a**) followed by hydrolysis of the ethyl ester protecting group gave the methyl acetamide derivative (**9**).

**Scheme 1.**
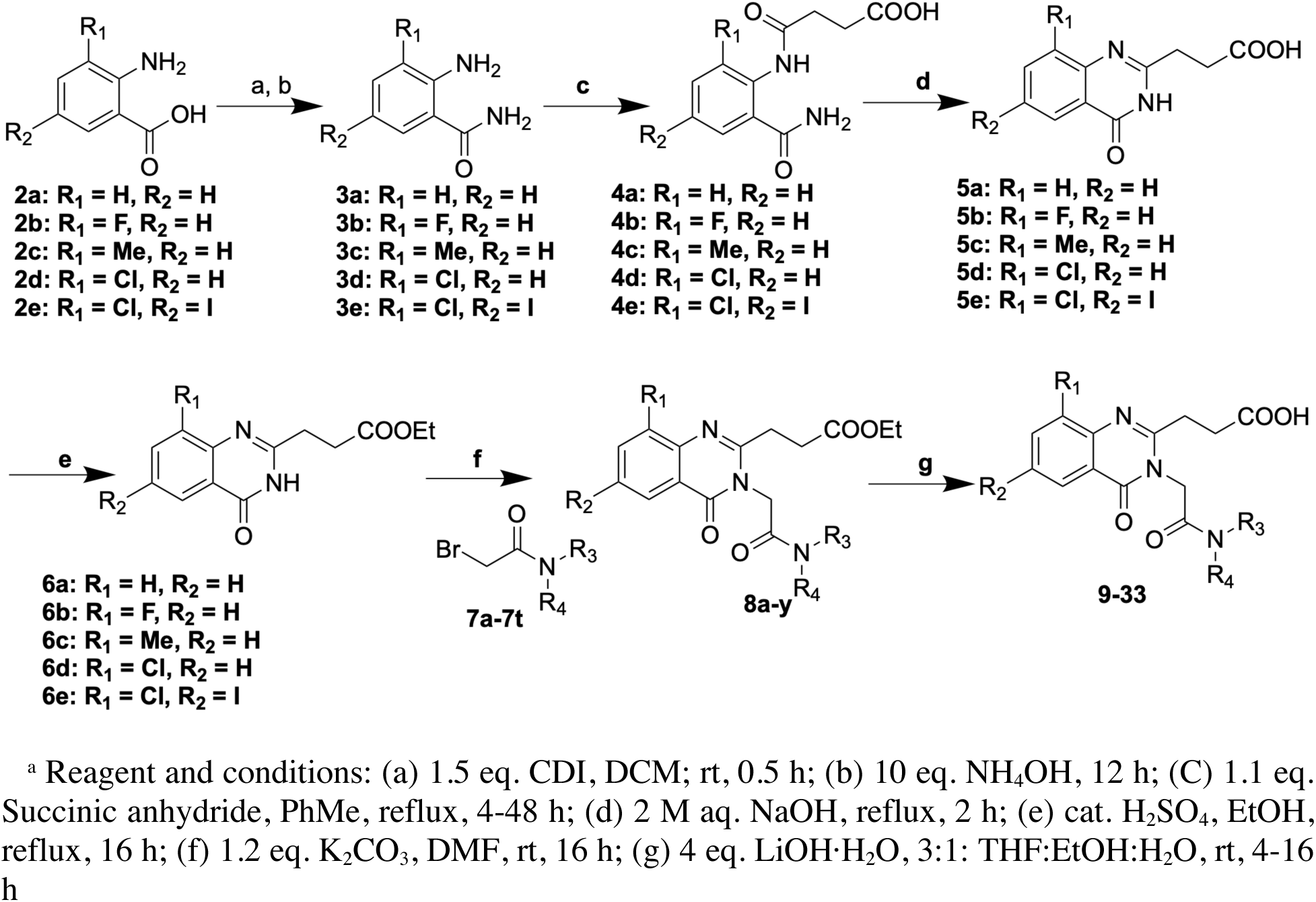
Synthesis of compounds **9 -32**^a^

Compared to **1**, the corresponding methylacetamide derivative (**9**) showed ∼4-fold loss of binding activity. However, we were able to obtain a 1.55 Å resolution (PDB: 8G43) co-crystal structure of **9** with HDAC6 UBD which revealed the potential to improve activity further (**Figure 2**). In the co-crystal structure, R1155 moves to open the side pocket. The carbonyl and NH groups of the amide of **9** make additional hydrogen bonds with R1155 and Y1189, respectively, compared to **1**, while maintaining the π-stacking of the quinazolinone core with R1155 and W1182 and hydrogen bond of the carboxylate with G1154 and Y1184. The co-crystal structure with **9** indicated that other larger amide groups could be extended into the side pocket to further improve activity.

**Figure 2.**
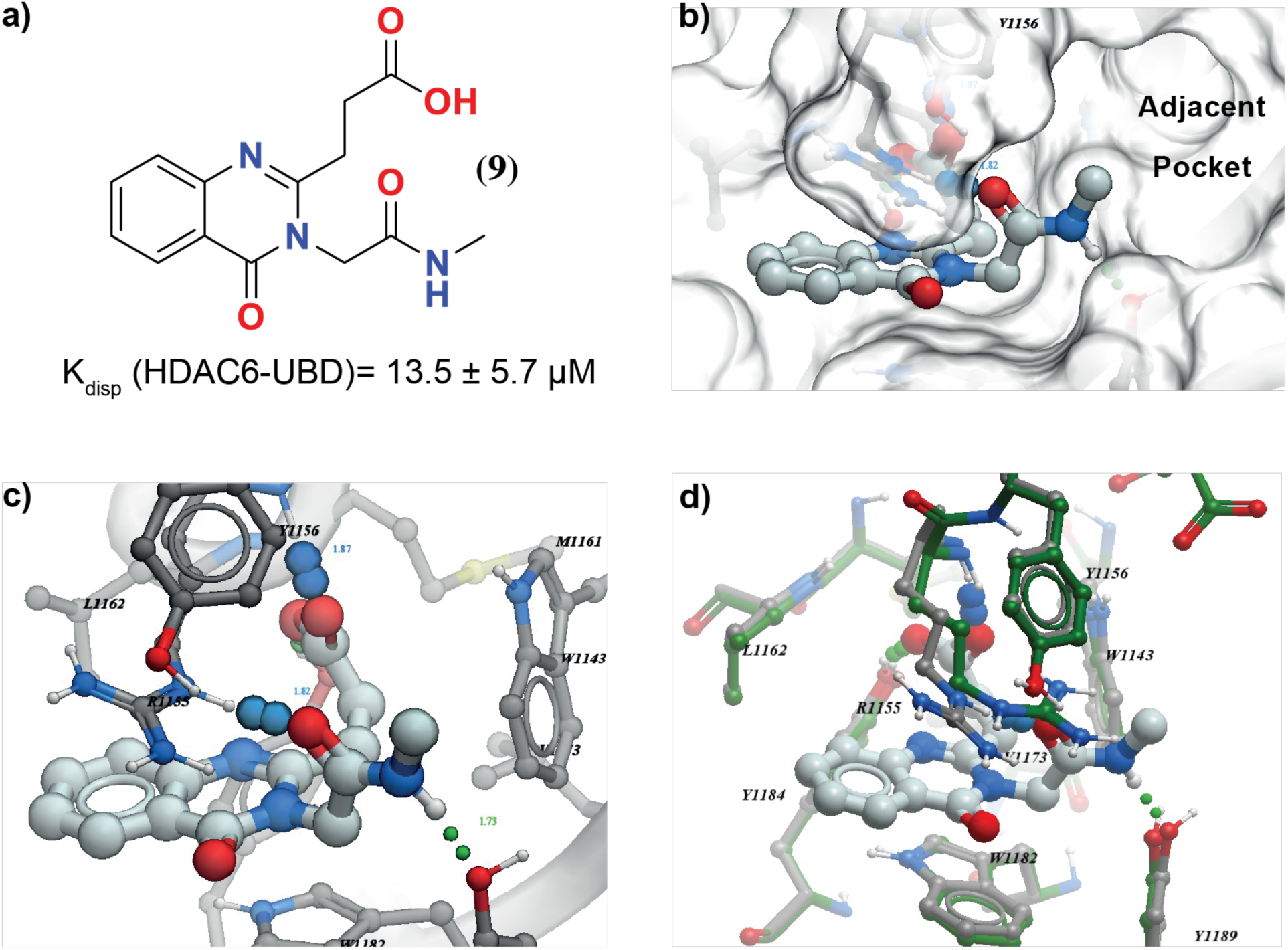
Structure and co-crystal structure of **9** with HDAC6 UBD. a) Structure of compound **9**. b) Space filled diagram of co-crystal structure of **9** with HDAC6 UBD showing adjacent pocket. c) Key hydrogen bond interactions formed in this complex. d) Overlay of the co-crystal structure of **9** (in grey) with **1** (in green) (for clarity **1** not shown), highlighting the movement of R1115 by arrow (in black).

To identify the optimal substitutions occupying the adjacent pocket and thus improve activity, we systematically explored an array of amides. These amide derivatives **10-22** were synthesized following Scheme 1 by using the corresponding amide derivatives of **7b-n** which were synthesized by simple amide coupling of commercially available amines with bromoacetyl bromide. The structure activity relationship of these amides is summarized in **Table 1**. While simple amides such as ethyl (**10**), cyclopropylmethyl (**11**), and tertiary butyl (**12**) amides are tolerated, they did not improve activity significantly. However, the larger lipophilic groups adamantyl (**13**), cyclohexylmethyl (**14**), and benzyl groups (**15**) improve the binding activity 3 –20 fold compared to methyl (**9)**, from K_disp_ = 13.5 μM (**9**) to K_disp_ = 5.1 μM, 2.0 μM, and 0.6 μM respectively.

**Table 1.**
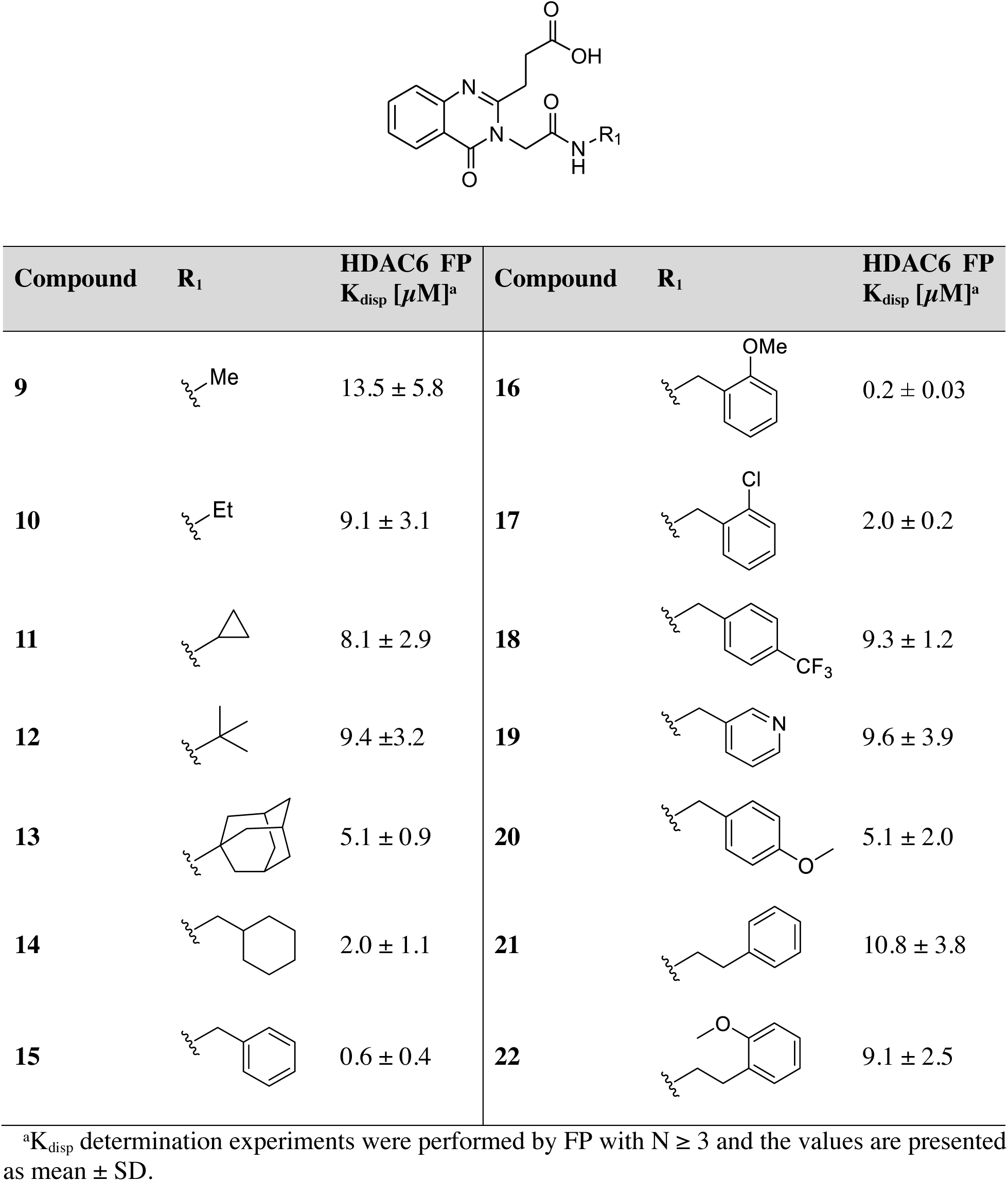
Structure and activity of amide substitutions (**9-22**)

| Compound | R<sub>1</sub> | HDAC6 FP<br>K<sub>disp</sub> [μM]<sup>a</sup> | Compound | R<sub>1</sub> | HDAC6 FP<br>K<sub>disp</sub> [μM]<sup>a</sup> |
| --- | --- | --- | --- | --- | --- |
| **9** |  | 13.5 ± 5.8 | **16** |  | 0.2 ± 0.03 |
| **10** |  | 9.1 ± 3.1 | **17** |  | 2.0 ± 0.2 |
| **11** |  | 8.1 ± 2.9 | **18** |  | 9.3 ± 1.2 |
| **12** |  | 9.4 ± 3.2 | **19** |  | 9.6 ± 3.9 |
| **13** |  | 5.1 ± 0.9 | **20** |  | 5.1 ± 2.0 |
| **14** |  | 2.0 ± 1.1 | **21** |  | 10.8 ± 3.8 |
| **15** |  | 0.6 ± 0.4 | **22** |  | 9.1 ± 2.5 |

<sup>a</sup>K<sub>disp</sub> determination experiments were performed by FP with N ≥ 3 and the values are presented as mean ± SD.

To confirm that the phenyl ring of **15** extends into the side pocket, we solved the crystal structure of HDAC6-UBD in complex with **15** to 1.55 Å resolution (PDB: 8G44) (**Figure 3**). As expected, the phenyl group of **15** occupies the side pocket and maintains the same interaction as **9** whilst making additional hydrophobic interactions with side pocket residues E1141 and I1177. The co-crystal structure with **15** also revealed that a hydrogen bond donor at the *ortho* position of the phenyl ring could pick up additional interaction with Y1189, and indeed, the *ortho*-OMe group (**16**) improved binding activity 3-fold compared to the phenyl analogue (**15**) while a Cl group at this position (**17**) led to 3-fold loss of activity. A CF_3_ group at the *para* position (**18**) was detrimental for binding. 3-Pyridyl (**19**) and *para*-OMe (**20**) analogues were designed to make additional interactions with D1178 or E1141 and S1175, respectively, but led to significant loss of activity compared to **15**. Extending the linker length of the amides (**21**, **22**) was not beneficial.

**Figure 3:**
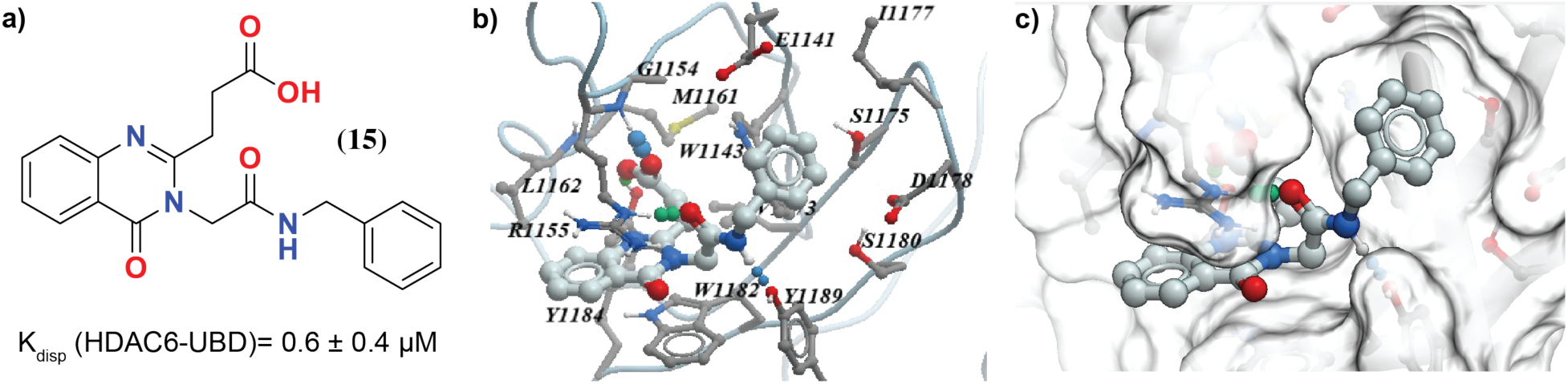
Structure and co-crystal structure of **15** with HDAC6 UBD. a) Structure of compound **15**. b) co-crystal structure of **15** with HDAC6 UBD showing key hydrogen bond interactions. c) space filled diagram showing a phenyl group occupying the adjacent pocket.

Based on the co-crystal structure of **15**, we envisioned that a simple alkyl or halo group at the 8-position of the quinazolinone ring could make hydrophobic interaction with the L1162 and thus improve activity further. Hydrophobic substitutions could also improve cell permeability so we also explored simple hydrophobic substitutions at the 6-position of the quinazolinone ring, which could point towards the adjacent pocket. To test this hypothesis, we synthesized analogues of **16** with substitutions at the 6-and/or 8-positions of the quinazolinone ring (**23-26**) according to Scheme 1, starting from appropriately substituted anthranilic acids (**2b-2e**). The structure activity relationship of substituted quinazolinone analogues is summarized in **Table 2**. While fluoro (**23**) and methyl (**24**) groups maintain activity, a chloro (**25**) group at the **8** position improves activity 2-fold compared to the unsubstituted quinazolinone analogues (**16**), from K_disp_ **=** 0.2 μM (**16**) to K_disp_ = 0.095 μM. An additional iodo group at the 6-position (**26**) was also tolerated without significant loss of activity.

**Table 2.**
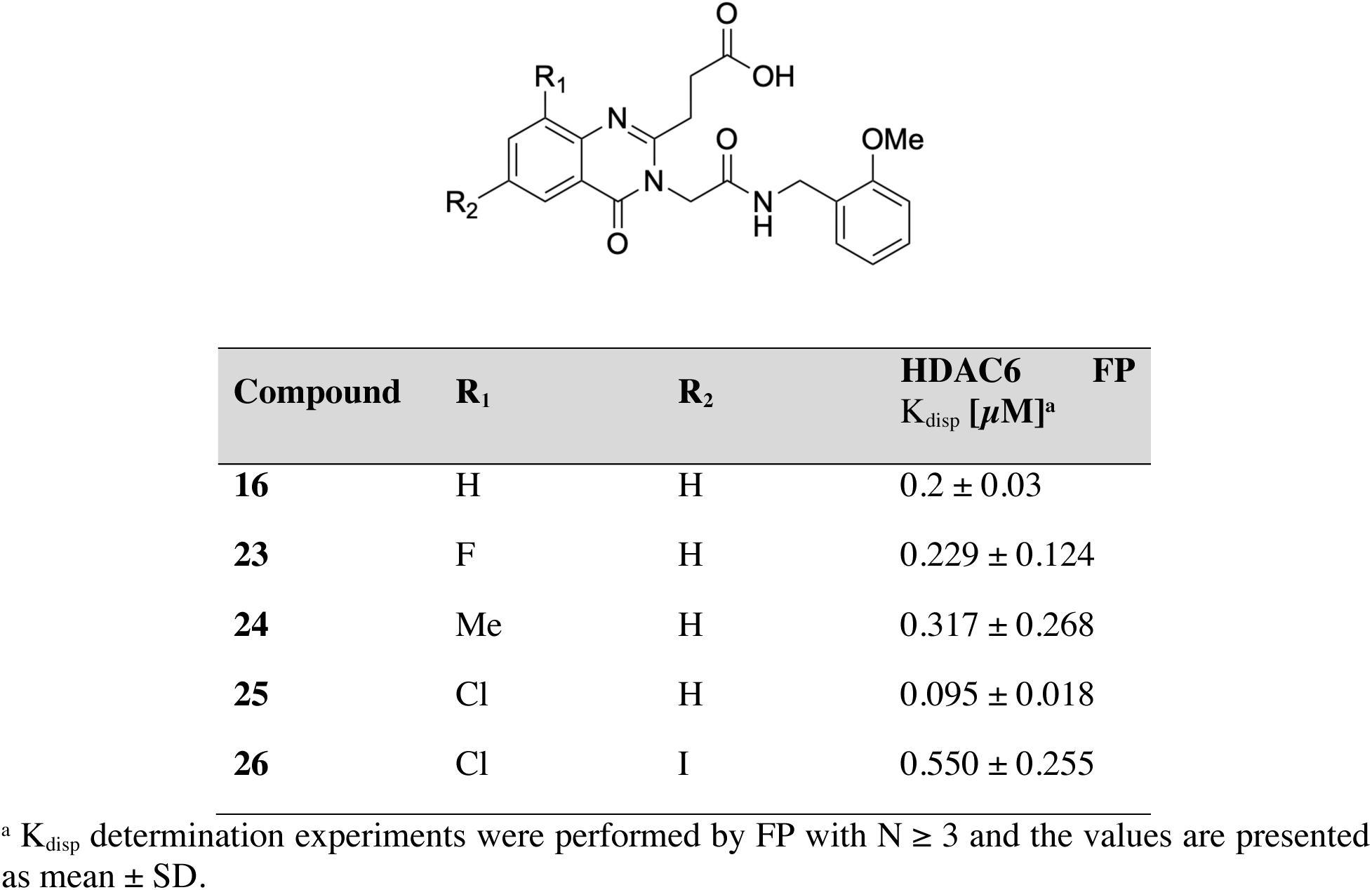
Structure and activity of substituted quinazolinones **16, 23 -26**

To identify opportunities to further improve the activity, we solved the crystal structure of HDAC6-UBD in complex with **25** to 1.55 Å resolution (PDB: 8G45) (**Figure 4**). As expected, the quinazolinone ring is sandwiched between R1155 and W1182 and maintains the same hydrogen bond interactions as **15** (**Figure 4**). However, the quinazolinone ring of **25** moves slightly away from L1161 to accommodate the chloro group without significant movement of the rest of the molecule or the sidechains of the binding site.

**Figure 4.**
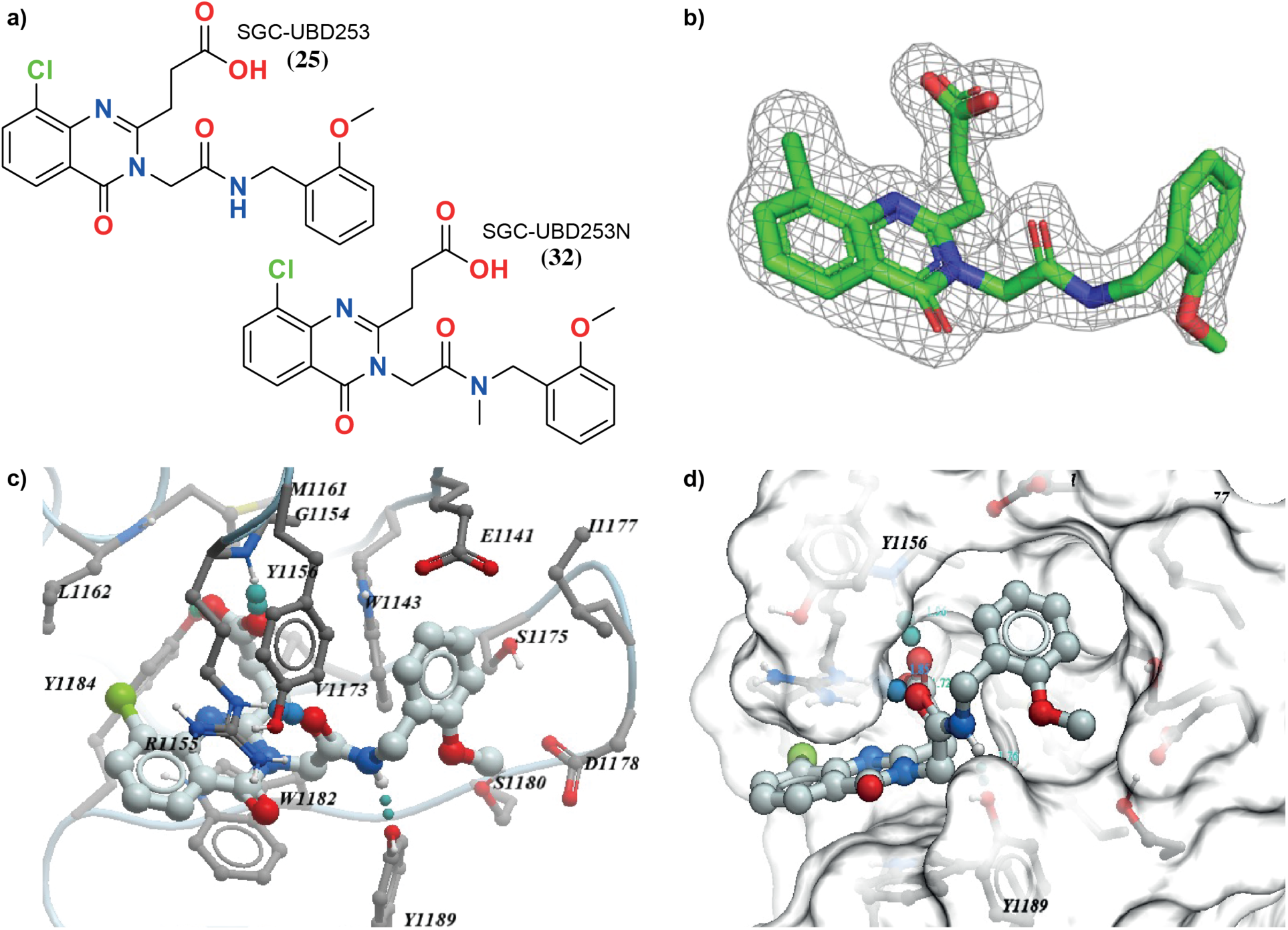
Structure and co-crystal structure of **25** with HDAC6 UBD. a) Structure of compound **25 (SGC-UBD253)** and negative control **32 (SGC-UBD253N)**, b) Omit map (σ2) of compound **25**, c) co-crystal structure of **25** with HDAC6 UBD showing key hydrogen bond interactions, d) space filled diagram showing phenyl group occupying the adjacent pocket.

With the optimal substitutions identified at the quinazolinone core, we wanted to identify the best amide group for binding activity and cellular activity. As amides with both hydrogen bond donors and with lipophilic substitutions were tolerated, we explored additional amide groups incorporating these features as summarized in **Table 3**.

**Table 3.**
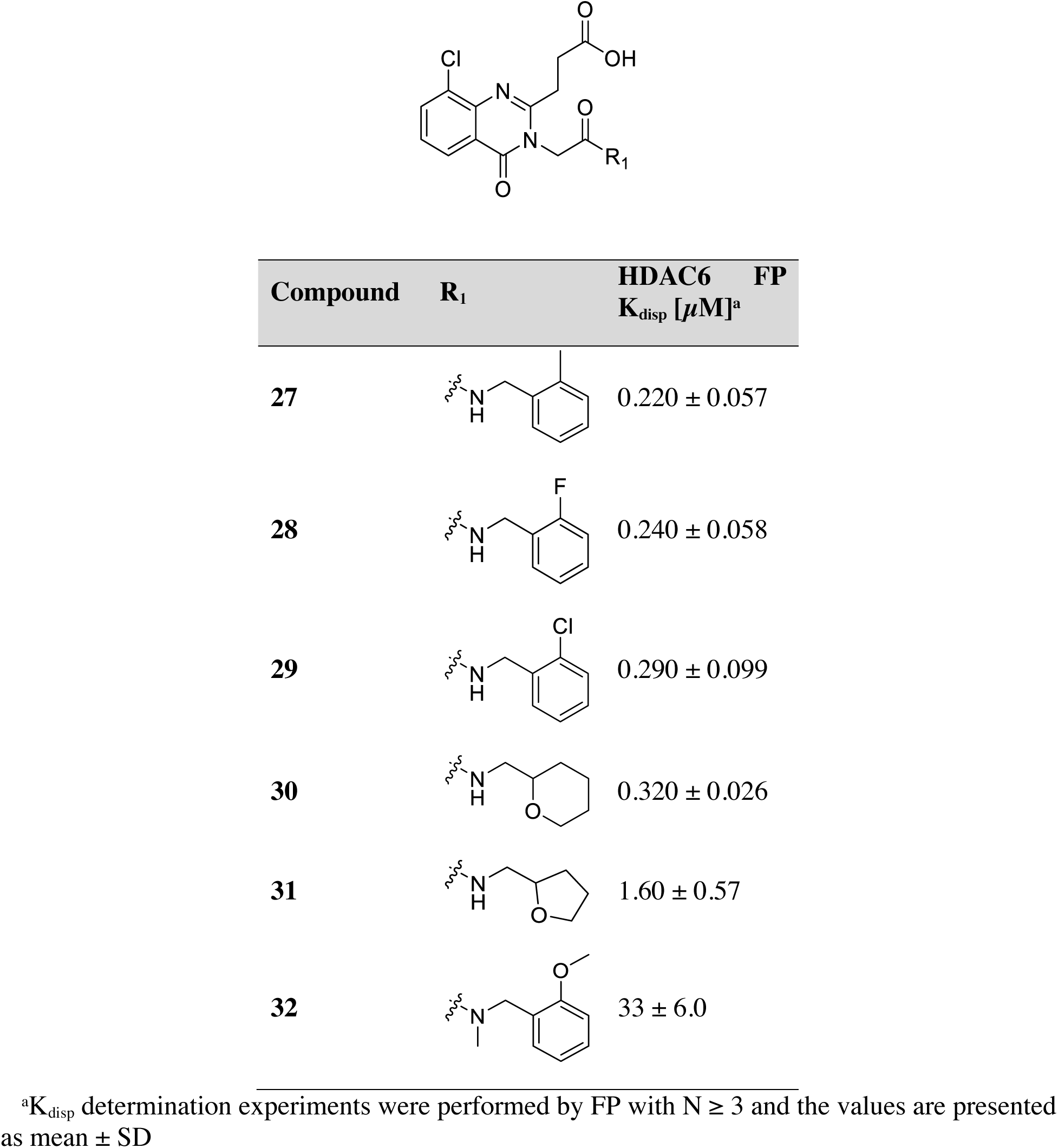
Structure and activity of amide substitutions (**27-32**)

To identify a close analog that may serve as negative control, we also synthesized **32** (**SGC-UBD253N**) by methylating the amide linker, which acts as a key hydrogen bond donor. As expected, **32** lost almost all the binding activity compared to the close analogue **25**, and is an excellent negative control for cellular experiments.

Furthermore, lipophilic substitutions at the *ortho* position, CH_3_ (**27**), F (**28**), and Cl (**29**) groups, showed comparable activity to the *ortho* OMe analogue (**25**). We explored tetrahydropyran (**30**) and tetrahydrofuran (**31**) analogues as both have oxygen atoms which could make hydrogen bond interactions with Y1189. While the tetrahydropyran (**30**) analogue showed a modest decrease in activity, the tetrahydrofuran (**31**) analogue showed a significant drop in activity (10-fold); probably because the tetrahydrofuran group may not occupy the pocket efficiently.

### IN VITRO CHARACTERIZATION OF **25** and **32**

**25** and **32** were selected as candidate probe and negative control compounds, both warranting further characterization by orthogonal biophysical assays (summarized in **Table 4** and **Figure S2**). Both were characterized by SPR and isothermal calorimetry (ITC). **25** binds potently to HDAC6-UBD with K_D_ values of 0.084 µM and 0.080 µM as measured by SPR and ITC, respectively. In contrast, **32** was shown to bind weakly with a K_D_ of 32 µM as determined by SPR. Binding parameters of **25** were also determined for full length HDAC6 using FP and SPR yielding K_disp_ and K_D_ values of 0.443 µM and 0.258 µM for these assays, respectively. As we determined with the C-terminal ubiquitin peptide substrate (**Table S1**), **25** binds more potently to the isolated HDAC6-UBD than the full-length protein.

**Table 4.**
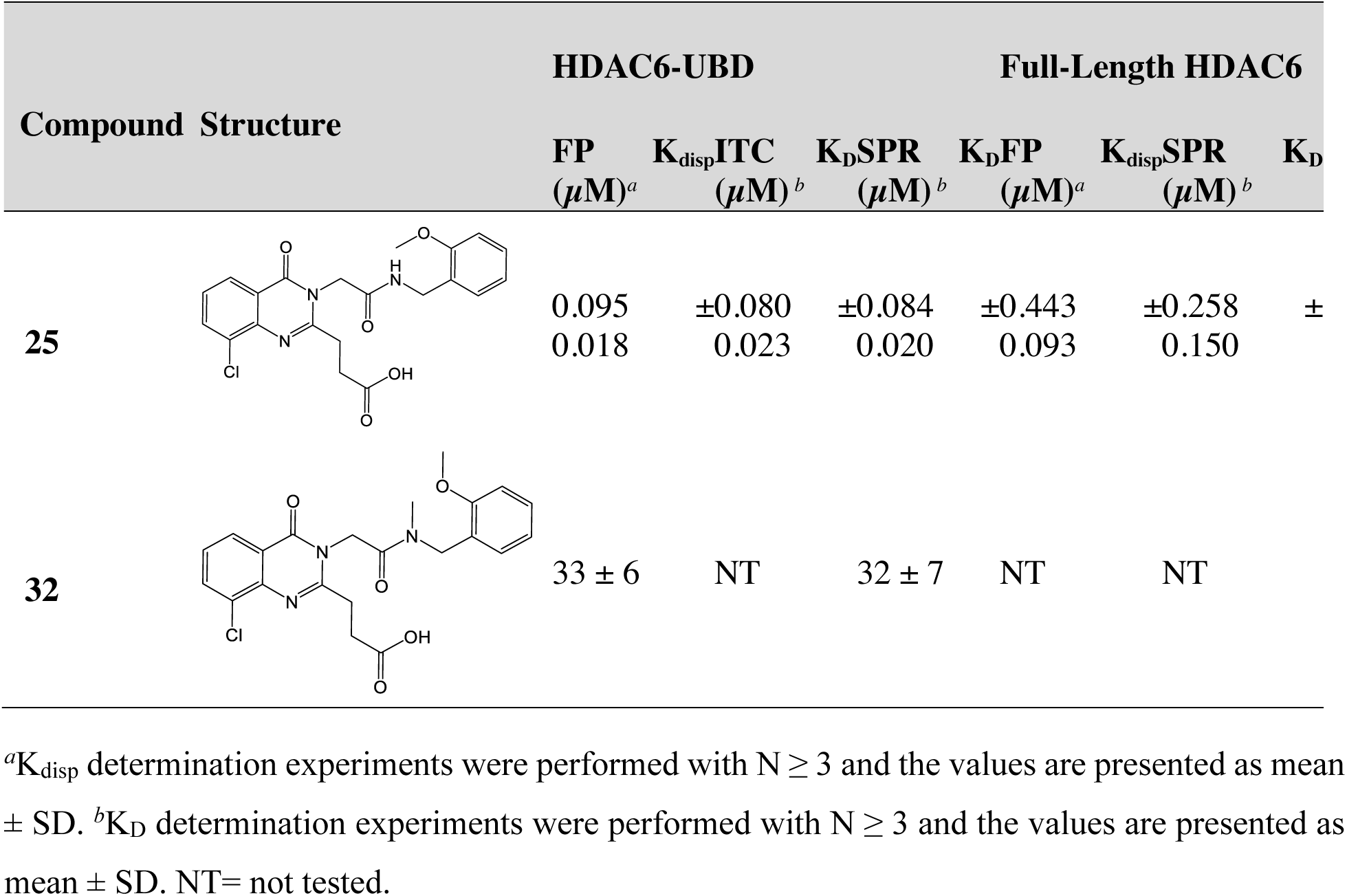
Summary of biophysical characterization of **25** and **32**

| Compound Structure | HDAC6-UBD |  |  | Full-Length HDAC6 |  |  |
| --- | --- | --- | --- | --- | --- | --- |
| | FP<br>($\mu\text{M}$)<sup>a</sup> | K<sub>disp</sub>ITC<br>($\mu\text{M}$)<sup>b</sup> | K<sub>D</sub>SPR<br>($\mu\text{M}$)<sup>b</sup> | K<sub>D</sub>FP<br>($\mu\text{M}$)<sup>a</sup> | K<sub>disp</sub>SPR<br>($\mu\text{M}$)<sup>b</sup> | K<sub>D</sub> |
| **25** | 0.095<br>0.018 | $\pm 0.080$<br>0.023 | $\pm 0.084$<br>0.020 | $\pm 0.443$<br>0.093 | $\pm 0.258$<br>0.150 | ± |
| **32** | $33 \pm 6$ | NT | $32 \pm 7$ | NT | NT | |

<sup>a</sup>K<sub>disp</sub> determination experiments were performed with N ≥ 3 and the values are presented as mean ± SD. <sup>b</sup>K<sub>D</sub> determination experiments were performed with N ≥ 3 and the values are presented as mean ± SD. NT= not tested.

Next, we tested the selectivity of **25** and **32** for HDAC6-UBD using a panel of UBD proteins. **25** was at least ∼50-fold selective for HDAC6 over other UBD proteins tested, with the exception of USP16, for which **25** is 15-fold selective as measured by SPR (**Table 5**). USP16 has the highest degree of conservation of binding pocket residues compared to HDAC6, with 53.3% sequence identity for residues within 5 Å of the bound ligand in the pocket (PDB ID: 8G45). Despite extensive chemistry efforts to improve selectivity of **25** for HDAC6 over USP16, we could not improve further while maintaining significant HDAC6 activity.

**Table 5.** Selectivity of **25** and **32** against a panel of UBDs by SPR

| UBD | $K_D$ (μM)<sup>a</sup><br>**25** | $K_D$ (μM)<sup>a</sup><br>**32** | Fold-selectivity of **25**<br>[UBD $K_D$ /HDAC6 $K_D$] |
| --- | --- | --- | --- |
| **USP3** | 11 ± 1.5 | 47 ± 4.4 | 130 |
| **USP5** | 4.5 ± 1.9 | 6.3 ± 2.1 | 53 |
| **USP13** | NB | NB | NC |
| **USP16** | 1.3 ± 0.04 | 4.9 ± 0.26 | 15 |
| **USP20** | NB | NB | NC |
| **USP33** | NB | NB | NC |
| **USP39** | NB | NB | NC |
| **USP49** | NB | NB | NC |
| **USP51** | NB | NB | NC |
| **BRAP** | NB | 92 ± 10 | NC |
| **HDAC6** | 0.084 ± 0.020 | 32 ± 7.3 |  |

<sup>a</sup> $K_D$ determination experiments were performed with $N \geq 3$ and the values are presented as mean ± SD. NB= $K_D$ greater than highest concentration of compound tested. NC= not calculated as no $K_D$ determined due to NB.

**25** was tested to assess possible inhibition of the deacetylase activity of HDAC6 using a Boc-Lys(TFA)-AMC substrate assay with this compound but no inhibition was observed (**Figure S3**). Similarly, inhibition of the ubiquitin peptidase activity of full-length USP proteins by **25** was assessed for USP3, USP5, USP16, and USP33 using a ubiquitin rhodamine substrate assay but no inhibition was observed, except for USP5. However, the IC_50_ in this *in vitro* assay of **25** for USP5 is 8 ± 1 µM so this is unlikely to have significant consequences for downstream use of this compound.

### CHARACTERIZATION OF **25** and **32** IN CELLS

We developed a nano-bioluminescence resonance energy transfer (NanoBRET) assay to quantitatively assess intracellular target engagement in real time in live cells (**Figure 5a**). This assay detects protein interactions, and their disruption with inhibitors, by measuring energy transfer from a bioluminescent protein donor to a fluorescent protein acceptor ^21^. ISG15 is a small ubiquitin-like modifier (SUMO) that is covalently attached to target proteins in a similar manner to ubiquitin. Importantly, both ISG15 and ubiquitin share the same C-terminal ‘LRLRGG’ motif ^22^, allowing ISG15 to also be recognized by HDAC6-UBD. For cellular NanoBRET assays, ISG15 was selected as the acceptor protein for the HDAC6 donor instead of ubiquitin because of the latter’s high cellular abundance and thus, high background signal in the assay. Using the HDAC6/ISG15 assay format, **25** was shown to significantly decrease the interaction between full-length HDAC6 and ISG15 compared to **32**, with EC_50_ values of 1.9 ± 0.6 µM and >30 μM, respectively, in HEK293T cells (**Figure 5b**). To validate the assay specificity, we identified a mutant HDAC6^R1155A, Y1184A^ which removes key hydrogen bond interactions with the terminal glycine of ubiquitin substrates resulting in decreased interaction with ISG15 *in vitro* (**Table S1**). Therefore, a donor construct bearing these mutations was used to define the baseline (binding deficient HDAC6) BRET signal in the assay (**Figure 5b**). Compound **25** was shown to disrupt the wildtype HDAC6/ISG15 BRET signal in a dose-dependent manner to the same level as HDAC6^R1155A, Y1184A^. To assess selectivity of **25** in cells, a NanoBRET assay was designed to assess USP16-ISG15 interaction. Similar to the HDAC6 NanoBRET assay, an ISG15-binding deficient mutant of USP16 (R84A/Y117A) was used to define the baseline of the assay. An EC_50_ of 20 ± 2.7 µM was determined using this assay in HEK293T cells, indicating approximately 10-fold selectivity of **25** for HDAC6 over USP16 under these conditions. The negative control candidate, **32**, was inactive. Analysis of the dose-dependent inhibition of both HDAC6 and USP16 suggests that using **25** at a concentration of 3 µM would limit off-target effects of USP16, whilst achieving significant antagonism of HDAC6.

**Figure 5.**
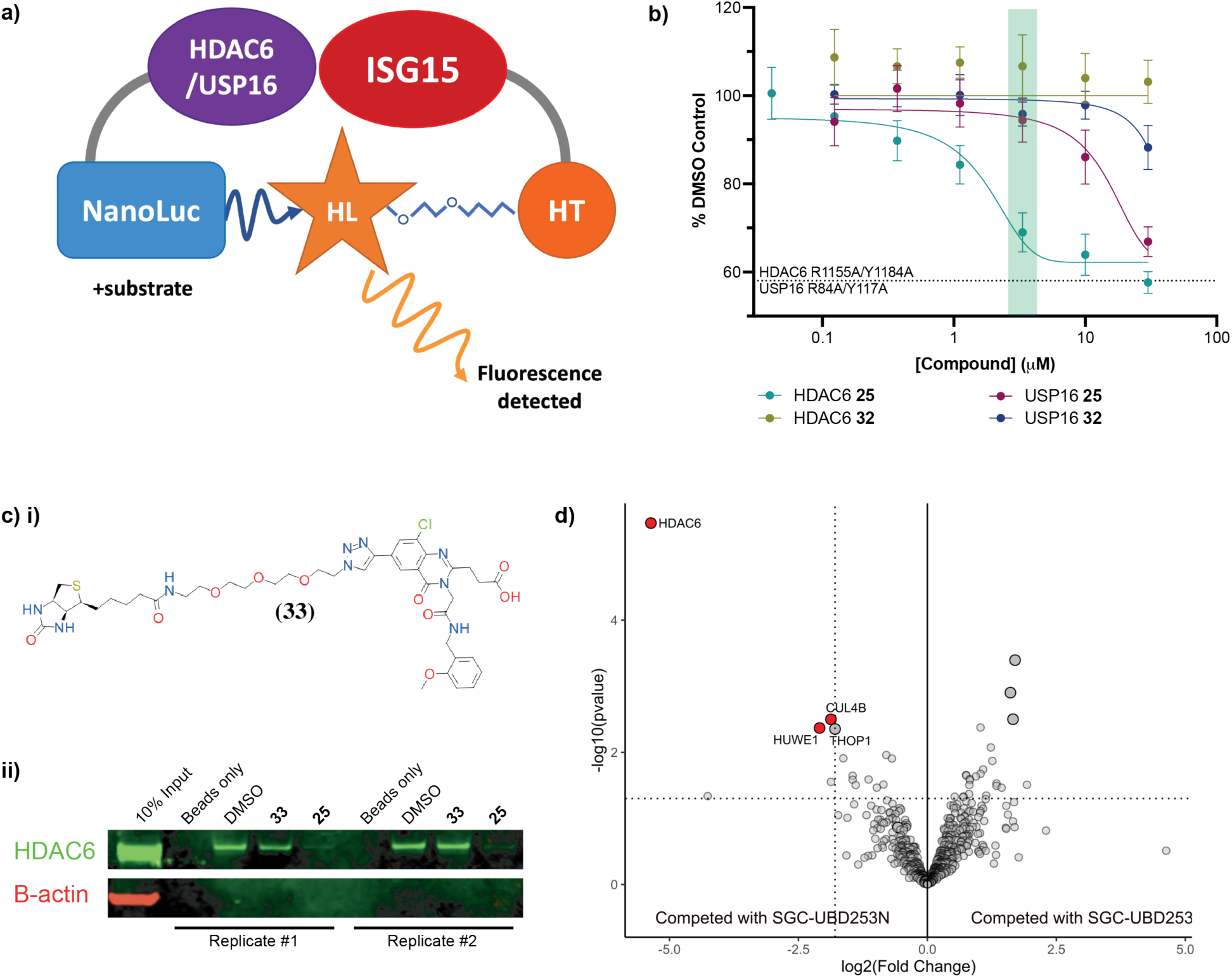
Assessment of cell-based target engagement of **25** and **32**. a) Schematic of NanoBRET assay which can measure the BRET from interaction between HDAC6 or USP16 tagged with NanoLuc donor and ISG15 tagged with acceptor HaloTag (HT) in the presence of HaloTag NanoBRET 618 Ligand (HL). b) **25** inhibited interaction of HDAC6 and ISG15 (EC_50_ =1.9 ± 0.6 µM) and USP16 and ISG15 (EC_50_=20 ± 2.7 µM) in HEK293T cells, whilst **32** was inactive. 3 µM is the recommended concentration (highlighted in green) to achieve selectivity for HDAC6 over USP16 for **25** in cells. Experimental triplicates of mean corrected NanoBRET ratios normalized to DMSO control are presented. c)i) Structure of compound **33** (the biotin-derivative of **25)**. ii) Western blot analysis of proteins pulled down from lysates by **33** after pretreatment with DMSO, of 10 µM of **32**, or **25**. Replicate experiments show that pre-treatment with **25**, but not **32**, inhibits HDAC6 pull-down with **33**. d) Selectivity profiling by label-free chemical proteomics in cytoplasmic fraction. The volcano plot is annotated with significantly enriched proteins where log2(Fold Change) < -2 and p-value ≤ 0.05 (n = 3 independent replicates). Data analysis was completed using the Bioconductor packages DEP ^24^. Parallel analysis using proDA ^25^ gave the output in **Supporting Data Files**.

To further characterize the selectivity of **25**, we synthesized **33** (**Figure 5ci**), a biotin-labeled derivative of **25**, for use as an affinity reagent for proteome-wide cellular target engagement profiling. The affinity of **33** for HDAC6-UBD was measured by FP and was found to be comparable to **25**. Two enrichment-based experiments were performed. First, we completed a competition assay in which HDAC6 was pulled down from HEK293T cell lysates treated with DMSO, **32**, or **25** (10 µM each). Treated lysates were incubated with **33** bound to streptavidin beads. After washing, the bound fraction was analyzed by western blot (**Figure 5cii**), revealing that pre-treatment with **25**, but not DMSO or **32**, inhibited pull down of HDAC6 from cells with **33**. Next, we performed chemical proteomics analysis of material pulled down from the cytoplasmic fraction of HEK293T cells by **33**-conjugated streptavidin beads. Pre-incubation with **25** prevented enrichment of HDAC6 by **33**, whereas pre-incubation with **32** did not affect the enrichment profile (**Figure 5d**). USP16 was not observed in the analysis from either pre-treatment condition. Since USP16 and HDAC6 are both localized to the cytoplasm ^2, 23^, this demonstrates that **25** has selectivity for endogenous HDAC6 over endogenous USP16.

## DISCUSSION AND CONCLUSIONS

We report the successful identification of a HDAC6-UBD chemical probe (**25**) (**SGC-UBD253**) and its negative control (**32**) (**SGC-UBD253N**) which we have thoroughly characterized with *in vitro* biophysical assays, crystal structures, and in functional cellular target engagement assays. **25** is a potent antagonist of HDAC6-UBD, active in cells, and shows good selectivity against other UBD domains, albeit with slight activity against the UBD domain of USP16 under some conditions. Additionally, we generated a biotin derivative of **25** which we validated as a useful tool for chemoproteomics experiments and demonstrated strong proteome-wide selectivity of **25** for HDAC6. We believe that the tools presented here will enable biological investigation of HDAC6-UBD and serve as a robust foundation for future applications such as targeted degradation agents and/or other proximity inducing agents.

## EXPERIMENTAL SECTION

### COMPOUND SYNTHESIS

#### General Considerations

Unless otherwise stated, all reactions were carried out under an inert atmosphere of dry argon or nitrogen utilizing glassware that was either oven (120 °C) or flame-dried. Workups and isolation of the products was conducted on the benchtop using standard techniques. Reactions were monitored using thin-layer chromatography (TLC) on SiliaPlate™ Silica Gel 60 F254 plates. Visualization of the plates was performed under UV light (254 nm) or using KMnO4 stains. Toluene was distilled over calcium hydride, and anhydrous N,N-dimethylformamide (DMF) was purchased from Fisher Sciences and used as received. Silica gel flash column chromatography was performed on Silicycle 230-400 mesh silica gel. Mono-and multidimensional NMR characterization data were collected at 298 K on a Varian Mercury 300, Varian Mercury 400, Bruker Avance II, Agilent 500, or a Varian 600. 1H NMR spectra were internally referenced to the residual solvent peak (CDCl_3_ = 7.26 ppm, DMSO-d6 = 2.50 ppm). 13C{1H} NMR spectra were internally referenced to the solvent peak (CDCl_3_ = 77.16 ppm, DMSO-d6 = 39.52 ppm). 19F NMR chemical shifts are reported in ppm with absolute reference to 1H. NMR data are reported as follows: chemical shift (δ ppm), multiplicity (s = singlet, d = doublet, t = triplet, q = quartet, m = multiplet, br = broad), coupling constant (Hz), integration. Coupling constants have been rounded to the nearest 0.05 Hz. The NMR spectra were recorded at the NMR facility of the Department of Chemistry at the University of Toronto. Infrared spectra were recorded on a Perkin-Elmer Spectrum 100 instrument equipped with a single-bounce diamond/ZnSe ATR accessory and are reported in wavenumber (cm-1) units. High resolution mass spectra (HRMS) were obtained on a micromass 70S-250 spectrometer (EI) or an ABI/SciexQStar Mass Spectrometer (ESI) or a JEOL AccuTOF model JMS-T1000LC mass spectrometer equipped with an IONICS® Direct Analysis in Real Time (DART) ion source at the Advanced Instrumentation for Molecular Structure (AIMS) facility of the Department of Chemistry at the University of Toronto. Analytical HPLC analyses were carried out on an Agilent 1100 series instrument equipped with a Phenomenex KINETEX® column (2.6 μm, C18, 50×4.6 mm). A linear gradient starting from 5% acetonitrile and 95% water (0.1% formic acid) to 95% acetonitrile and 5% water (0.1% formic acid) over 4 minutes followed by 5 minutes of elution at 95% acetonitrile and 5% water (0.1% formic acid) was employed. Flow rate was 1 mL/min, and UV detection was set to 254 nm and 214 nm. HPLC analyses were conducted at room temperature. All compounds submitted for testing were at ≥95% purity (HPLC) unless otherwise stated.

#### General Procedure 1 (Synthesis of α-Bromoamide Derivatives 7b-7t)

The aldehyde derivative (1 eq.) was added dropwise to a suspension of the hydroxylamine·HCl salt in absolute ethanol (1.5 M) and stirred at room temperature for 3-16 h (depending on the ketone or aldehyde). Upon consumption of the starting material, concentrated HCl (4 eq.) was added to the reaction. The solution was then cooled to 0 °C and zinc powder was added portion-wise. Following complete addition of the zinc powder, the reaction was warmed to room temperature and stirred for 10 minutes. A 2:1 solution of 6 M NaOH and NH_4_OH (1.07 M, relative to starting material) was added, resulting in the formation of a white precipitate. The suspension was filtered through a pad of celite. The white solids were washed three times with DCM. The filtrate was collected and extracted three times with DCM, dried over MgSO_4_, filtered, and concentrated *in vacuo*. The amine was used in the next step without further purification. Bromoacetyl bromide (1.2 eq.) was added dropwise at 0 °C to a solution of the amine (1 eq.) and K_2_CO_3_ (1.2 eq.) in DCM (0.4 M). The solution was warmed to room temperature and stirred for 4-12 h. The reaction was carefully quenched by dropwise addition of water. The aqueous layer was extracted three times with DCM. The organic layers were combined, dried over MgSO_4_, filtered, and concentrated *in vacuo*. The resultant α-bromoamides were used without further purification.

#### General Procedure 2 (Synthesis of Quinazolinone Derivatives: 9-32)

**3a-3e**: The anthranilic acid derivative (1 eq.) and CDI (1.5 eq.) were added to a round-bottomed flask connected to an oil bubbler. The reagents were dissolved in DCM (0.4 M) and stirred at room temperature for 0.5 h. The round-bottom was fitted with an addition funnel, and NH_4_OH (10 eq.) was added dropwise. The reaction was stirred at room temperature for 12 h. The reaction was concentrated *in vacuo*. The residue was dissolved in EtOAc, washed twice with 1 M HCl, once with an aqueous solution of NaHCO_3_, once with water, and once with brine. The organic layer was dried over MgSO_4_, filtered, and concentrated *in vacuo*. The resulting anthranilamide derivatives were used in the next step without further purification.

**4a-4e**: In a flask equipped with a reflux condenser, succinic anhydride (1.1 eq.) and the anthranilamide **(3a-e)** (1 eq.) were dissolved in toluene and stirred at reflux for 4-48 h. The reaction was cooled to room temperature and filtered. The white precipitate was washed with water, Et_2_O, and a small amount of absolute ethanol. The product was dried and used without further purification in the next step.

**5a-5e**: In a flask equipped with a reflux condenser, the intermediate bis-amide (**4a-e**) was dissolved in 2 M NaOH (0.2 M). The resulting solution was stirred at a reflux and then cooled to room temperature or treated with 2 M K_2_CO_3_ at room temperature for 2 hours. The solution was carefully acidified to pH 4-6 using concentrated HCl while stirring. The white precipitate was filtered and washed with water and dried. The resulting quinazolinone was used in the next step without further purification.

**6a-6e**: The acid (**5a-e**) was dissolved in EtOH (0.1 M) in a flask equipped with a reflux condenser. A catalytic amount of H_2_SO_4_ (1 drop/mmol) was added, and the solution was stirred at a reflux for 16 h. The reaction was cooled to room temperature and diluted with water, and then cooled to -40 °C in a freezer. The resulting white solid was filtered, washed with a saturated solution of NaHCO_3_, and then water, and dried. The product was used in the next step without further purification.

**8a-8t**: NaH (2 eq.) was added to a solution of the secondary amide (**6a-e**) (2 eq.) in THF (0.4 M) at 0°C. The solution was stirred at 0 °C until bubbling had ceased. α-Bromoamide derivatives (**7a-s**) (2 eq.) was added dropwise at 0 °C. The solution was warmed to room temperature and stirred for 12 h. The reaction was quenched with water and diluted with EtOAc. The aqueous layer was extracted three times with EtOAc. The organics were combined, dried over MgSO_4_, filtered, and concentrated *in vacuo*. The resulting tertiary amide was used in the next step without further purification.

**9-32**: In a flask containing the ester (**8a-t**) (1 eq.) and LiOH·H_2_O (4 eq.) was added a solution of 3:1:1 THF:H_2_O:EtOH (0.05 M). Upon complete consumption of the starting material, the solution was neutralized to pH 6-7 using 1 M HCl. The organics were removed *in vacuo*, and the resulting solid was filtered, washed with water, acetone, and then Et_2_O. In some cases the material was further purified by flash column chromatography (1:1 acetone:PhMe with 0.1% acetic acid) or by other means (specified for each compound).

**3-(3-(2-(methylamino)-2-oxoethyl)-4-oxo-3,4-dihydroquinazolin-2-yl)propanoic acid (9):** ^1^H NMR (DMSO-*d*_6_, 500 MHz): δ 12.18 (s, 1H), 8.23 (q, *J* = 4.6 Hz, 1H), 8.09 (dd, *J* = 7.9, 1.6 Hz, 1H), 7.81 (ddd, *J* = 8.4, 7.1, 1.6 Hz, 1H), 7.58 (dd, *J* = 8.2, 1.1 Hz, 1H), 7.50 (ddd, *J* = 8.1, 7.1, 1.1 Hz, 1H), 2.97 (t, *J* = 6.8 Hz, 2H), 4.78 (s, 2H), 2.75 (t, *J* = 6.8 Hz, 2H), 2.64 (d, *J* = 4.6 Hz, 3H). ^13^C{^1^H} NMR (DMSO-*d*_6_, 125 MHz): δ 173.7, 166.9, 161.2, 156.3, 146.7, 134.4, 126.8, 126.4, 126.2, 119.7, 45.2, 30.0, 28.7, 25.7. IR (neat): 3298, 2936, 1675, 1655, 1597, 1588, 1391, 1236, 1185, 974, 905, 765, 705. HRMS: ESI+, calc. for C_14_H_16_N_3_O_4_ 290.11408 [M+H] ^+^, found 290.11398.

**3-(3-(2-(ethylamino)-2-oxoethyl)-4-oxo-3,4-dihydroquinazolin-2-yl)propanoic acid (10):** ^1^H NMR (DMSO-*d*_6_, 500 MHz): δ 12.17 (s, 1H), 8.31 (t, *J* = 5.5 Hz, 1H), 8.09 (dd, *J* = 8.0, 1.5 Hz, 1H), 7.81 (ddd, *J* = 8.4, 7.1, 1.6 Hz, 1H), 7.58 (dd, *J* = 8.2, 1.0 Hz, 1H), 7.50 (ddd, *J* = 8.2, 7.1, 1.2 Hz, 1H), 4.78 (s, 2H), 3.12 (qd, *J* = 7.2, 5.4 Hz, 2H), 2.97 (t, *J* = 6.8 Hz, 2H), 2.75 (t, *J* = 6.8 Hz, 2H), 1.05 (t, *J* = 7.2 Hz, 3H). ^13^C{^1^H} NMR (DMSO-*d*_6_, 125 MHz): δ 174.1, 166.6, 161.6, 156.8, 147.1, 134.9, 127.2, 126.9, 126.7, 120.2, 45.6, 34.1, 30.5, 29.2, 15.0. IR (neat): 3300, 2927, 1653, 1597, 1555, 1393, 1184, 942, 766, 708, 690. HRMS: ESI+, calc. for C_15_H_18_N_3_O_4_ 304.12973 [M+H] ^+^, found 304.12995.

**3-(3-(2-(cyclopropylamino)-2-oxoethyl)-4-oxo-3,4-dihydroquinazolin-2-yl)propanoic acid (11):** ^1^H NMR (DMSO-*d*_6_, 500 MHz): δ 12.18 (s, 1H), 8.41 (d, *J* = 4.1 Hz, 1H), 8.09 (ddd, *J* = 7.9, 1.6, 0.6 Hz, 1H), 7.81 (ddd, *J* = 8.6, 7.1, 1.6 Hz, 1H), 7.58 (dt, *J* = 8.0, 0.9 Hz, 1H), 7.50 (ddd, *J* = 8.2, 7.1, 1.2 Hz, 1H), 4.74 (s, 2H), 2.96 (t, *J* = 6.8 Hz, 2H), 2.75 (t, *J* = 6.8 Hz, 2H), 2.62 – 2.71 (m, 1H), 0.61 – 0.67 (m, 2H), 0.42 – 0.47 (m, 2H). ^13^C{^1^H} NMR (DMSO-*d*_6_, 125 MHz): δ 173.6, 167.5, 161.1, 156.3, 146.6, 134.4, 126.8, 126.4, 126.2, 119.7, 45.1, 30.0, 28.7, 22.4, 5.6. IR (neat): 3283, 3083, 2960, 1648, 1654, 1598, 1557, 1397, 1272, 1182, 775, 708. HRMS: ESI+, calc. for C_16_H_18_N_3_O_4_ 316.12973 [M+H] ^+^, found 316.13027.

**3-(3-(2-(tert-butylamino)-2-oxoethyl)-4-oxo-3,4-dihydroquinazolin-2-yl)propanoic acid (12):** ^1^H NMR (DMSO-*d*_6_, 500 MHz): δ 12.16 (s, 1H), 8.09 (ddd, *J* = 7.9, 1.6, 0.6 Hz, 1H), 8.00 (s, 1H), 7.80 (ddd, *J* = 8.2, 7.1, 1.6 Hz, 1H), 7.58 (ddd, *J* = 8.2, 1.1, 0.5 Hz, 1H), 7.50 (ddd, *J* = 8.1, 7.1, 1.2 Hz, 1H), 4.76 (s, 2H), 2.94 (t, *J* = 6.8 Hz, 2H), 2.75 (t, *J* = 6.8 Hz, 2H), 1.27 (s, 9H). ^13^C{^1^H} NMR (DMSO-*d*_6_, 125 MHz): δ 173.6, 165.6, 161.1, 156.4, 146.7, 134.4, 126.8, 126.4, 126.2, 119.6, 50.5, 45.2, 30.0, 28.7, 28.4. IR (neat): 3473, 3298, 3097, 2981, 2927, 1722, 1636, 1592, 1567, 1394, 1364, 1222, 1183, 780. HRMS: ESI+, calc. for C_17_H_22_N_3_O_4_ 332.16103 [M+H]^+^, found 332.16063.

**3-(3-(2-(adamantan-1-yl)amino)-2-oxoethyl)-4-oxo-3,4-dihydroquinazolin-2-yl)propanoic acid (13):** ^1^H-NMR (DMSO-*d*_6_, 500 MHz): δ 12.14 (s, 1H), 8.09 (dd, *J* = 8.0, 1.5 Hz, 1H), 7.88 (s, 1H), 1.61 (s, 6H), 7.80 (ddd, *J* = 8.5, 7.1, 1.6 Hz, 1H), 7.58 (d, *J* = 8.1 Hz, 1H), 7.45 – 7.53 (m, 1H), 4.76 (s, 2H), 2.93 (t, *J* = 6.8 Hz, 2H), 2.75 (t, *J* = 6.8 Hz, 2H), 2.01 (s, 3H), 1.94 (s, 6H). ^13^C{^1^H}-NMR (DMSO-*d*_6_, 125 MHz): δ 173.6, 165.3, 161.1, 156.4, 146.6, 134.4, 126.7, 126.4, 126.2, 119.6, 51.2, 45.2, 40.9, 30.0, 28.8, 28.7. IR (neat): 3309, 2910, 2853, 2663, 1600, 1547, 1343, 1278, 1245, 1182, 780. HRMS: ESI+, calc. for C_23_H_28_N_3_O_4_ 410.20798 [M+H] ^+^, found 410.20843.

**3-(3-(2-((cyclohexylmethyl)amino)-2-oxoethyl)-4-oxo-3,4-dihydroquinazolin-2-yl)propanoic acid (14):** ^1^H NMR (DMSO-*d_6_*, 500 MHz): δ 12.18 (s, 1H), 8.28 (t, *J* = 5.9 Hz, 1H), 8.09 (dd, *J* = 8.0, 1.5 Hz, 1H), 7.80 (ddd, *J* = 8.5, 7.1, 1.6 Hz, 1H), 7.58 (dd, *J* = 8.2, 0.9 Hz, 1H), 7.49 (ddd, *J* = 8.2, 7.1, 1.2 Hz, 1H), 4.80 (s, 2H), 3.04 – 2.89 (m, 2H), 2.75 (t, *J* = 6.9 Hz, 2H), 1.74 – 1.64 (m, 4H), 1.63 – 1.57 (m, 1H), 1.49 – 1.31 (m, 1H), 1.26 – 1.03 (m, 3H), 0.95 – 0.79 (m, 2H). ^13^C{^1^H} NMR (DMSO-*d_6_,* 125 MHz): δ 173.6, 166.4, 161.2, 156.3, 146.7, 134.4, 126.8, 126.4, 126.2, 119.7, 45.1, 45.1, 37.5, 30.4, 30.0, 28.8, 26.0, 25.4. HRMS: ESI+, calc. for C_20_H_26_N_3_O_4_ 372.19233 [M+H] ^+^, found 372.19233.

**3-(3-(2-(benzylamino)-2-oxoethyl)-4-oxo-3,4-dihydroquinazolin-2-yl)propanoic acid (15):** ^1^H NMR (DMSO-*d*_6_, 500 MHz): δ 8.90 (t, *J* = 5.8 Hz, 1H), 8.11 (dd, *J* = 8.0, 1.5 Hz, 1H), 7.81 (ddd, *J* = 8.5, 7.2, 1.6 Hz, 1H), 7.60 (dd, *J* = 8.2, 1.0 Hz, 1H), 7.51 (ddd, *J* = 8.1, 7.1, 1.1 Hz, 1H), 4.89 (s, 2H), 7.21 – 7.40 (m, 5H), 4.34 (d, *J* = 5.9 Hz, 2H), 3.01 (t, *J* = 6.9 Hz, 2H), 2.76 (t, *J* = 6.8 Hz, 2H). The signal of the COOH group is extremely broad. ^13^C{^1^H} NMR (DMSO-*d*_6_, 125 MHz): δ 173.6, 166.6, 161.2, 156.4, 146.5, 139.0, 134.5, 128.3, 127.2, 126.9, 126.7, 126.5, 126.2, 119.7, 45.3, 42.3, 30.1, 28.8. IR (neat): 3285, 3065, 2940, 1652, 1597, 1552, 1391, 1248, 1179, 971, 774, 744, 695. HRMS: ESI+, calc. for C_20_H_20_N_3_O_4_ 366.14538 [M+H] ^+^, found 366.14496.

**3-(3-(2-((2-methoxybenzyl)amino)-2-oxoethyl)-4-oxo-3,4-dihydroquinazolin-2-yl)propanoic acid (16):** ^1^H NMR (DMSO-*d_6_,* 500 MHz): δ 12.20 (s, 1H), 8.67 (t, *J* = 5.8 Hz, 1H), 8.11 (dd, *J* = 8.0, 1.6 Hz, 1H), 7.81 (ddd, *J* = 8.5, 7.1, 1.6 Hz, 1H), 7.65 – 7.57 (m, 1H), 7.53 – 7.45 (m, 1H), 7.34 – 7.18 (m, 2H), 6.98 (d, *J* = 8.0 Hz, 1H), 6.93 (t, *J* = 7.3 Hz, 1H), 4.89 (s, 2H), 4.29 (d, *J* = 5.7 Hz, 2H), 3.80 (s, 3H), 3.01 (t, *J* = 6.8 Hz, 2H), 2.76 (t, *J* = 6.8 Hz, 2H). ^13^C{^1^H} NMR (DMSO-*d_6_,* 125 MHz): δ 173.7, 166.6, 161.2, 156.7, 156.3, 146.7, 134.5, 128.2, 127.9, 126.8, 126.5, 126.3, 126.2, 119.7, 110.5, 55.3, 45.2, 37.5, 30.1, 28.8. HRMS: ESI+, calc. for C_21_H_22_N_3_O_5_ 396.15595 [M+H] ^+^, found 396.15592.

**3-(3-(2-((2-chlorobenzyl)amino)-2-oxoethyl)-4-oxo-3,4-dihydroquinazolin-2-yl)propanoic acid (17):** ^1^H NMR (DMSO-*d_6_*, 500 MHz): δ 12.22 (s, 1H), 8.88 (t, *J* = 5.8 Hz, 1H), 8.11 (dd, *J* = 7.9, 1.5 Hz, 1H), 7.81 (ddd, *J* = 8.5, 7.2, 1.6 Hz, 1H), 7.59 (dd, *J* = 8.3, 1.0 Hz, 1H), 7.50 (ddd, *J* = 8.2, 7.1, 1.2 Hz, 1H), 7.43 (ddd, *J* = 12.0, 7.6, 1.7 Hz, 2H), 7.38 – 7.25 (m, 2H), 4.92 (s, 2H), 4.40 (d, *J* = 5.7 Hz, 2H), 3.02 (t, *J* = 6.8 Hz, 2H), 2.76 (t, *J* = 6.8 Hz, 2H). ^13^C{^1^H} NMR (DMSO-*d_6_*, 125 MHz): δ 173.7, 166.9, 161.2, 156.2, 146.7, 135.9, 134.5, 132.1, 129.2, 128.9, 128.8, 127.2, 126.8, 126.5, 126.2, 119.7, 45.3, 40.3, 30.1, 28.8. HRMS: ESI+, calc. for C_20_H_19_ClN_3_O_4_ 400.10641 [M+H] ^+^, found 400.10649.

**3-(4-oxo-3-(2-oxo-2-((4-(trifluoromethyl)benzyl)amino)ethyl)-3,4-dihydroquinazolin-2-yl)propanoic acid (18**): ^1^H NMR (DMSO-*d_6_*, 400 MHz): δ 9.19 (t, *J* = 6.0 Hz, 1H), 8.10 (dd, *J* = 8.1, 1.1 Hz, 1H), 7.80 (ddd, *J* = 8.5, 7.1, 1.6 Hz, 1H), 7.70 (d, *J* = 8.0 Hz, 2H), 7.60 (dt, *J* = 8.2, 1.0 Hz, 1H), 7.56 – 7.45 (m, 3H), 4.43 (d, *J* = 5.8 Hz, 2H), 2.94 (t, *J* = 7.1 Hz, 2H), 2.60 (t, *J* = 7.1 Hz, 2H). ^13^C{^1^H} NMR (DMSO-*d_6_,* 100 MHz,): δ 174.3, 167.2, 161.4, 157.8, 147.0, 144.1, 134.3, 127.8, 127.6, 127.3, 126.7, 126.2, 125.2 (1, *J* = 3.6 Hz), 119.7, 45.7, 41.9, 33.1, 30.4. ^19^F NMR (DMSO-d_6_, 377 MHz) δ -60.8. IR (neat): 3281, 2930, 1657, 1597, 1548, 1421, 1387, 1327, 1251, 1166, 1122, 1069, 778, 703, 648. HRMS: DART, calc. for C_21_H_19_F_3_N_3_O_4_ 434.13277 [M+H] ^+^, found 434.13209.

**3-(4-oxo-3-(2-oxo-2-((pyridin-3-ylmethyl)amino)ethyl)-3,4-dihydroquinazolin-2-yl)propanoic acid (19):** ^1^H NMR (DMSO-*d_6_,* 400 MHz): δ 12.19 (s, 1H), 8.90 (t, *J* = 5.9 Hz, 1H), 8.52 (s, 1H), 8.47 (d, *J* = 4.7 Hz, 1H), 8.10 (dd, *J* = 8.0, 1.5 Hz, 1H), 7.81 (ddd, *J* = 8.4, 7.1, 1.6 Hz, 1H), 7.69 (d, *J* = 7.9 Hz, 1H), 7.59 (d, *J* = 8.1 Hz, 1H), 7.50 (td, *J* = 7.6, 7.1, 1.2 Hz, 1H), 7.36 (dd, *J* = 7.8, 4.7 Hz, 1H), 4.88 (s, 2H), 4.37 (d, *J* = 5.8 Hz, 2H), 3.00 (t, *J* = 6.8 Hz, 2H), 2.76 (t, *J* = 6.8 Hz, 2H). ^13^C{^1^H} NMR (DMSO-*d_6_,* 101 MHz): δ 173.7, 166.9, 161.2, 156.2, 148.7, 148.2, 146.7, 135.0, 134.5, 134.5, 126.8, 126.5, 126.2, 123.5, 119.7, 45.4, 40.0, 30.0, 28.8. HRMS: DART, calc. for C_20_H_18_ClFN_3_O_4_ 418.09644 [M+H] ^+^, found 418.09634.

**3-(3-(2-((4-methoxybenzyl)amino)-2-oxoethyl)-4-oxo-3,4-dihydroquinazolin-2-yl)propanoic acid (20):** ^1^H NMR (DMSO-*d_6_*, 500 MHz): δ 12.21 (s, 1H), 8.77 (t, *J* = 5.9 Hz, 1H), 8.10 (d, *J* = 7.9 Hz, 1H), 7.81 (t, *J* = 7.7 Hz, 1H), 7.59 (d, *J* = 8.1 Hz, 1H), 7.50 (t, *J* = 7.6 Hz, 1H), 7.21 (d, *J* = 8.2 Hz, 2H), 6.89 (d, *J* = 8.2 Hz, 2H), 4.86 (s, 2H), 4.26 (d, *J* = 5.9 Hz, 2H), 3.73 (s, 3H), 3.00 (t, *J* = 6.9 Hz, 2H), 2.76 (t, *J* = 6.8 Hz, 2H). ^13^C{^1^H} NMR (DMSO-*d_6_*, 125 MHz): δ 173.7, 166.5, 161.2, 158.3, 156.3, 146.7, 134.5, 130.9, 128.61 126.8, 126.5, 126.3, 119.8, 113.8, 55.1, 45.3, 41.9, 30.1, 28.8. HRMS: ESI+, calc. for C_21_H_22_N_3_O_5_ 396.15595 [M+H] ^+^, found 396.15549.

**3-(4-oxo-3-(2-oxo-2-(phenethylamino)ethyl)-3,4-dihydroquinazolin-2-yl)propanoic acid (21):** ^1^H NMR (DMSO-*d_6_*, 500 MHz): δ 12.20 (s, 1H), 8.41 (t, *J* = 5.6 Hz, 1H), 8.10 (dd, *J* = 8.0, 1.5 Hz, 1H), 7.81 (ddd, *J* = 8.5, 7.1, 1.6 Hz, 1H), 7.58 (dd, *J* = 8.2, 1.0 Hz, 1H), 7.50 (ddd, *J* = 8.1, 7.2, 1.2 Hz, 1H), 7.36 – 7.27 (m, 2H), 7.25 – 7.15 (m, 3H), 4.77 (s, 2H), 3.40 – 3.25 (m, 3H), 2.91 (t, *J* = 6.8 Hz, 2H), 2.74 (td, *J* = 7.1, 3.2 Hz, 4H). ^13^C{^1^H} NMR (DMSO-*d_6_*, 125 MHz): δ 173.6, 166.4, 161.1, 156.2, 146.6, 139.3, 134.4, 128.7, 128.3, 126.8, 126.4, 126.2, 126.1, 119.7, 45.1, 40.4, 35.0, 30.0, 28.7. HRMS: ESI+, calc. for C_21_H_22_N_3_O_4_ 380.16103 [M+H] ^+^, found 380.16087.

**3-(3-(2-((2-methoxyphenethyl)amino)-2-oxoethyl)-4-oxo-3,4-dihydroquinazolin-2-yl)propanoic acid (22):** ^1^H NMR (DMSO-*d_6_*, 500 MHz): δ 12.19 (s, 1H), 8.38 (t, *J* = 5.7 Hz, 1H), 8.09 (dd, *J* = 8.0, 1.5 Hz, 1H), 7.81 (ddd, *J* = 8.4, 7.1, 1.6 Hz, 1H), 7.64 – 7.55 (m, 1H), 7.50 (ddd, *J* = 8.1, 7.1, 1.1 Hz, 1H), 7.20 (ddd, *J* = 8.3, 7.4, 1.8 Hz, 1H), 7.13 (dd, *J* = 7.4, 1.8 Hz, 1H), 6.95 (dd, *J* = 8.3, 1.0 Hz, 1H), 6.87 (td, *J* = 7.4, 1.1 Hz, 1H), 4.77 (s, 2H), 3.78 (s, 3H), 3.32 – 3.26 (m, 2H), 2.92 (t, *J* = 6.8 Hz, 2H), 2.74 (dt, *J* = 10.3, 6.8 Hz, 4H). ^13^C{^1^H} NMR (DMSO-*d_6_*, 125 MHz): δ 173.7, 166.3, 161.1, 157.2, 156.2, 146.7, 134.4, 130.1, 127.6, 126.9, 126.8, 126.4, 126.2, 120.3, 119.7, 110.7, 55.3, 45.1, 38.9, 30.0, 29.9, 28.7. HRMS: DART, calc. for C_22_H_24_N_3_O_5_ 410.17160 [M+H] ^+^, found 410.17235.

**3-(8-fluoro-3-(2-((2-methoxybenzyl)amino)-2-oxoethyl)-4-oxo-3,4-dihydroquinazolin-2-yl)propanoic acid (23):** ^1^H NMR (DMSO-*d_6_*, 500 MHz): δ 8.68 (t, *J* = 5.8 Hz, 1H), 7.91 (d, *J* = 8.0 Hz, 1H), 7.68 (t, *J* = 9.2 Hz, 1H), 7.48 (td, *J* = 8.0, 4.5 Hz, 1H), 7.25 (ddd, *J* = 10.2, 7.4, 2.4 Hz, 2H), 6.98 (d, *J* = 8.1 Hz, 1H), 6.92 (t, *J* = 7.4 Hz, 1H), 4.90 (s, 2H), 4.30 (d, *J* = 5.7 Hz, 2H), 3.80 (s, 3H), 3.03 (t, *J* = 6.9 Hz, 2H), 2.77 (t, *J* = 6.8 Hz, 2H). ^13^C{^1^H} NMR (DMSO-*d_6_*, 125 MHz): δ 173.6, 166.4, 160.4 (d, *J* = 3.2 Hz), 157.2, 157.0, 156.7, 155.0, 136.0 (d, *J* = 11.6 Hz), 128.3, 127.9, 126.8 (d, *J* = 7.5 Hz), 126.2, 121.8, 120.2, 110.5, 55.4, 55.3, 45.5, 37.6, 30.0, 29.1. ^19^F NMR (DMSO-*d_6_*, 377 MHz): δ -126.28. HRMS: DART, calc. for C_21_H_21_FN_3_O_5_ 414.14352 [M+H] ^+^, found 414.14687.

**3-(3-(2-((2-methoxybenzyl)amino)-2-oxoethyl)-8-methyl-4-oxo-3,4-dihydroquinazolin-2-yl)propanoic acid (24):** ^1^H NMR (DMSO-*d_6_*, 500 MHz): δ 8.69 (t, *J* = 5.8 Hz, 1H), 7.92 (dd, *J* = 8.0, 1.6 Hz, 1H), 7.64 (d, *J* = 7.2 Hz, 1H), 7.36 (t, *J* = 7.6 Hz, 1H), 7.25 (td, *J* = 7.3, 6.8, 1.9 Hz, 2H), 6.97 (d, *J* = 8.1 Hz, 1H), 6.93 (t, *J* = 7.4 Hz, 1H), 4.89 (s, 2H), 4.29 (d, *J* = 5.7 Hz, 2H), 3.80 (s, 3H), 2.99 (t, *J* = 6.6 Hz, 2H), 2.73 (t, *J* = 6.6 Hz, 2H), 2.52 (s, 3H). ^13^C{^1^H} NMR (DMSO-*d_6_*, 125 MHz): δ 174.1, 166.7, 161.5, 156.7, 155.4, 145.1, 135.0, 134.6, 128.2, 127.9, 126.3, 125.9, 120.2, 119.6, 110.5, 55.3, 55.3, 45.2, 37.5, 30.8, 29.4, 16.7. HRMS: DART, calc. for C_22_H_24_N_3_O_5_ 410.17160 [M+H] ^+^, found 410.17144.

**3-(8-chloro-3-(2-((2-methoxybenzyl)amino)-2-oxoethyl)-4-oxo-3,4-dihydroquinazolin-2-yl)propanoic acid (25):** ^1^H NMR (DMSO-*d*_6_, 500 MHz): δ 12.20 (br s, 1H), 8.68 (t, *J* = 5.8 Hz, 1H), 8.06 (dd, *J* = 8.0, 1.4 Hz, 1H), 7.97 (dd, *J* = 7.7, 1.4 Hz, 1H), 7.48 (t, *J* = 7.9 Hz, 1H), 7.17 – 7.32 (m, 2H), 6.98 (d, *J* = 7.7 Hz, 1H), 6.92 (td, *J* = 7.4, 1.0 Hz, 1H), 4.89 (s, 2H), 4.29 (d, *J* = 5.7 Hz, 2H), 3.80 (s, 3H), 3.04 (t, *J* = 6.7 Hz, 2H), 2.80 (t, *J* = 6.7 Hz, 2H). ^13^C{^1^H} NMR (DMSO-*d*_6_, 125 MHz): δ 173.5, 166.3, 160.7, 157.3, 156.7, 143.1, 134.5, 130.4, 128.3, 127.9, 126.9, 126.2, 125.4, 121.4, 120.2, 110.5, 55.3, 45.5, 37.5, 29.8, 29.1. IR (neat): 3294, 2977, 1746, 1722, 1690, 1658, 1610, 1434, 1401, 1373, 1233, 1261, 756. HRMS: ESI+, calc for C_21_H_21_ClN_3_O_5_ 430.11697 [M+H]^+^, found 430.11749.

**3-(8-chloro-6-iodo-3-(2-((2-methoxybenzyl)amino)-2-oxoethyl)-4-oxo-3,4-dihydroquinazolin-2-yl)propanoic acid (26):** ^1^H NMR (DMSO-*d_6_*, 500 MHz): δ 1H NMR (400 MHz, DMSO-d6) δ 12.21 (s, 1H), 8.69 (t, J = 5.8 Hz, 1H), 8.34 (d, J = 1.9 Hz, 1H), 8.32 (d, J = 1.9 Hz, 1H), 7.30 – 7.25 (m, 1H), 7.25 – 7.21 (m, 1H), 7.00 (d, J = 8.1 Hz, 1H), 6.97 – 6.90 (m, 1H), 4.89 (s, 2H), 4.30 (d, J = 5.7 Hz, 2H), 3.81 (s, 3H), 3.04 (t, J = 6.7 Hz, 2H), 2.79 (t, J = 6.7 Hz, 2H). HRMS: ESI calc. for C_21_H_19_ClIN_3_O_5_ 556.01307 [M+H] +, found 556.01.

**3-(8-chloro-3-(2-((2-methylbenzyl)amino)-2-oxoethyl)-4-oxo-3,4-dihydroquinazolin-2-yl)propanoic acid (27):** ^1^H NMR (CD_3_OD, 500 MHz): δ 8.09 (dd, *J* = 7.9, 1.4 Hz, 1H), 7.85 (dd, *J* = 7.7, 1.4 Hz, 1H), 7.39 (t, *J* = 7.9 Hz, 1H), 7.34 – 7.23 (m, 1H), 7.20 – 7.05 (m, 3H), 4.98 (s, 2H), 4.42 (s, 2H), 3.08 (t, *J* = 7.0 Hz, 2H), 2.88 (t, *J* = 7.0 Hz, 2H), 2.33 (s, 3H). ^13^C{^1^H} NMR (DMSO-*d_6_,* 125 MHz,): δ 179.1, 169.0, 163.5, 158.9, 145.3, 137.4, 137.0, 135.7, 132.9, 131.3, 129.3, 128.5, 127.6, 127.1, 126.3, 122.9, 47.0, 42.6, 33.7, 31.9, 19.1. IR (neat): 3359, 3283, 3080, 2957, 1687, 1645, 1580, 1422, 1386, 1327, 1246, 1225, 1170, 1003, 994, 981, 770, 761, 683, 451 HRMS: ESI+, calc. for C_21_H_21_ClN_3_O_4_ 414.12206 [M+H] ^+^, found 414.12249.

**3-(8-chloro-3-(2-((2-fluorobenzyl)amino)-2-oxoethyl)-4-oxo-3,4-dihydroquinazolin-2-yl)propanoic acid (28):** ^1^H NMR (DMSO-*d_6_*, 500 MHz): δ 8.87 (t, *J* = 5.8 Hz, 1H), 8.05 (dd, *J* = 8.0, 1.4 Hz, 1H), 7.95 (dd, *J* = 7.7, 1.4 Hz, 1H), 7.47 (t, *J* = 7.9 Hz, 1H), 7.38 (td, *J* = 7.8, 1.9 Hz, 1H), 7.32 (tdd, *J* = 7.5, 5.3, 1.8 Hz, 1H), 7.21 – 7.12 (m, 2H), 4.89 (s, 2H), 4.38 (d, *J* = 5.8 Hz, 2H), 3.03 (t, *J* = 6.7 Hz, 2H), 2.80 (t, *J* = 6.7 Hz, 2H). HRMS: DART, calc. for C_20_H_18_ClFN_3_O_4_ 418.09644 [M+H] ^+^, found 418.09634.

**3-(8-chloro-3-(2-((2-chlorobenzyl)amino)-2-oxoethyl)-4-oxo-3,4-dihydroquinazolin-2-yl)propanoic acid (29):** ^1^H NMR (DMSO-*d_6_*, 500 MHz): δ 8.98 (t, *J* = 5.9 Hz, 1H), 8.05 (d, *J* = 8.0 Hz, 1H), 7.94 (d, *J* = 7.7 Hz, 1H), 7.44 (dt, *J* = 15.3, 7.7 Hz, 3H), 7.31 (dt, *J* = 25.1, 7.4 Hz, 2H), 4.94 (s, 2H), 4.40 (d, *J* = 5.7 Hz, 2H), 3.02 (t, *J* = 7.0 Hz, 2H), 2.75 (t, *J* = 6.9 Hz, 2H). ^13^C{^1^H} NMR (DMSO-*d_6_,* 125 MHz,): δ 174.1, 166.8, 160.8, 157.9, 143.2, 135.9, 134.5, 132.0, 130.4, 129.1, 128.9, 128.7, 127.3, 126.8, 125.4, 121.4, 45.7, 31.2, 29.8. HRMS: DART, calc. for C_20_H_18_Cl_2_N_3_O_4_ 434.06744 [M+H] ^+^, found 434.06703.

**3-(8-chloro-4-oxo-3-(2-oxo-2-(((tetrahydro-2H-pyran-2-yl)methyl)amino)ethyl)-3,4-dihydroquinazolin-2-yl)propanoic acid (30):** ^1^H NMR (DMSO-*d_6_*, 500 MHz): δ 12.17 (s, 1H), 8.43 (t, *J* = 5.8 Hz, 1H), 8.04 (dd, *J* = 8.0, 1.4 Hz, 1H), 7.96 (dd, *J* = 7.8, 1.4 Hz, 1H), 7.47 (t, *J* = 7.9 Hz, 1H), 4.83 (s, 2H), 3.91 – 3.84 (m, 1H), 3.38 – 3.27 (m, 2H), 3.17 (ddd, *J* = 13.6, 6.0, 4.5 Hz, 1H), 3.08 (ddd, *J* = 13.6, 7.0, 5.6 Hz, 1H), 3.00 (t, *J* = 6.8 Hz, 2H), 2.79 (t, *J* = 6.7 Hz, 2H), 1.79 – 1.73 (m, 1H), 1.58 – 1.51 (m, 1H), 1.49 – 1.38 (m, 3H), 1.21 – 1.09 (m, 1H). ^13^C{^1^H} NMR (DMSO-*d_6_,* 125 MHz,): δ 173.6, 166.3, 160.7, 157.4, 143.1, 134.5, 130.4, 126.9, 125.4, 121.4, 75.8, 67.3, 45.3, 44.0, 29.8, 29.0, 28.9, 25.6, 22.6. IR (neat): 3445, 3285, 2946, 1726, 1654, 1597, 1573, 1447, 1201, 1165, 1096, 1047, 993, 925, 906, 768, 686. HRMS: ESI+, calc for C_19_H_23_ClN_3_O_5_ [408.13207] [M+H] ^+^, found [408.13234].

**3-(8-chloro-4-oxo-3-(2-oxo-2-(((tetrahydrofuran-2-yl)methyl)amino)ethyl)-3,4-dihydroquinazolin-2-yl)propanoic acid (31):** ^1^H NMR (DMSO-*d_6_*, 500 MHz): δ 12.18 – 12.15 (m, 1H), 8.45 (t, *J* = 5.8 Hz, 1H), 8.04 (dd, *J* = 8.0, 1.4 Hz, 1H), 7.95 (dd, *J* = 7.8, 1.4 Hz, 1H), 7.47 (t, *J* = 7.9 Hz, 1H), 4.83 (s, 2H), 3.85 (qd, *J* = 6.6, 5.0 Hz, 1H), 3.77 (ddd, *J* = 8.2, 7.0, 6.1 Hz, 1H), 3.66 – 3.58 (m, 1H), 3.25 – 3.12 (m, 2H), 3.00 (t, *J* = 6.7 Hz, 2H), 2.79 (t, *J* = 6.7 Hz, 2H), 1.93 – 1.72 (m, 3H), 1.50 (ddt, *J* = 12.5, 9.0, 7.3 Hz, 1H). ^13^C{^1^H} NMR (DMSO-*d_6_,* 125 MHz,): δ 173.6, 166.4, 160.7, 157.4, 143.1, 134.5, 130.4, 126.8, 125.4, 121.4, 77.1, 67.2, 45.3, 42.9, 29.8, 29.0, 28.5, 25.2. IR (neat): 3289, 3087, 2938, 1705, 1680, 1652, 1596, 1556, 1441, 1328, 1169, 1138, 1069, 984, 796, 764. HRMS: ESI+, calc for C_18_H_21_ClN_3_O_5_ [394.11642] [M+H]^+^, found [394.11661].

**3-(8-chloro-3-(2-((2-methoxybenzyl)(methyl)amino)-2-oxoethyl)-4-oxo-3,4-dihydroquinazolin-2-yl)propanoic acid (32):** ^1^H NMR (DMSO-*d_6_*, 500 MHz) (Hindered rotation): δ 8.04 (dd, 1H), 7.95 (d, 1H), 7.46 (t, 1H), 7.33 (t, 0.5H), 7.24 (t, 1H), 7.07 (t, 1H), 7.01 (t, 1H), 6.92 (t, 0.5H), 5.19 (d, 2H), 4.63 - 4.50 (2xs, 2H), 3.88 - 3.79 (2xs, 3H), 3.10-2.95 (s,t,t 3.5H), 2.77 -2.48 (2xt, 3.5H). HRMS: ESI+, calc. for C_22_H_23_ClN_3_O_5_ 444.13262 [M+H]^+^, found 444.13319.

### PROTEIN PURIFICATION

HDAC6^1109–1213^ with an N-terminal His-tag and a thrombin protease site, ISG15^81–157^ with an N-terminal His-tag and a TEV protease site, were expressed in *Escherichia coli* BL21 (DE3) codon plus cells from a pET28a-LIC and pET28-MHL vector, respectively. HDAC6^1109–1215^, HDAC6^1108^– _1215 R1155A_Y1184A, USP31-131, USP5171-290, USP1625-185, USP201-141, USP3329-134, USP3984-194, USP491-115,_ USP51^176–305^ and BRAP^304–390^ encoding an N-terminal AviTag for biotinylation and a C-terminal His6 tag were expressed in BirA cells from p28BIOH-LIC vectors, while USP13^183–307^ with an N-terminal His-tag and TEV protease cleavage site, and a C-terminal biotinylation sequence was expressed in BirA cells from a pNICBIO2 vector. All cells, apart from cells expressing ISG15^81–, 157^ and USP5^1–835^ were grown in M9 minimal media in the presence of 50 µM ZnSO_4_, 50 µg/mL of kanamycin and 30 µg/mL of chloramphenicol to an OD_600_ of 0.8 and induced by isopropyl-1-thio-D-galactopyranoside (IPTG) for a final concentration of 0.5 mM; cultures were incubated overnight for 18 hours at 15 ⁰C. ISG15^81–157^ and USP5^1–835^ were grown in Terrific broth (TB) media (Sigma Aldrich) with the same conditions. All p28BIOH-LIC or pNICBIO2 expressing cells were also supplemented with 1x of 100x biotin stock (10 mM; 2.442 mg/mL).

HDAC6^1–1215^, USP3^1–520^, USP16^1–823^, and USP33^c36–825^ were overexpressed in Sf9 cells where cultures were grown in HyQ SFX Insect Serum Free Medium (Fisher Scientific) to a density of 4 x 10^6^ cells/mL and infected with 10 mL of P3 viral stock per 1 L of cell culture. Cell culture medium was collected after 4 days of incubation in a shaker at 27 ⁰C.

Cells were harvested by centrifugation at 4,000-6,000 RPM at 10 ⁰C and cell pellets were lysed by sonification in lysis buffer 50 mM Tris pH 8, 150 mM NaCl, 1 mM TCEP (for proteins: _HDAC61109-1213, ISG1581-157, HDAC61109-1215, HDAC61108-1215 R1155A_Y1184A, USP31-131, USP5171-290,_ USP13^183–307^, USP16^25–185^, USP20^1–141^, USP33^29–134^, USP39^84–194^, USP49^1–115^, USP51^176–305^ and BRAP^304–390^, USP16^1–823^), 50 mM Tris pH 8, 500 mM NaCl, 2.5% glycerol (v/v), 1 mM TCEP (for proteins: USP3^1–520^, USP33^c36–825^), 50 mM Tris pH 8, 150 mM NaCl, 1 mM TCEP, 5% glycerol (v/v) (for protein: USP5^1–835^) supplemented with 50 µL benzonase, 1 mM protease inhibitor (PMSF, benzamidine) and 0.5% NP-40 (only for sf9 expression). The crude extract was centrifuged at 14,000 RPM for 1 hour at 10 ⁰C. The clarified lysate was incubated with nickel-nitrilotriacetic acid (Ni-NTA) agarose resin (Qiagen) for 1 hour with agitation. The protein bound resin was washed in wash buffer 1 (lysis buffer with no additives), then wash buffer 2 (wash buffer 1 supplemented with 15 mM imidazole), and finally bound proteins were eluted with elution buffer (wash buffer 1 supplemented with 300 mM imidazole) and monitored with Bradford reagent.

HDAC6^1109–1213^ was dialyzed overnight in 2 L of wash buffer 1 with 100 U of thrombin. Dialyzed sample was supplemented with 2 mM CaCl_2_ and treated daily with 100 U of thrombin, until protein cleavage was complete, as assessed by SDS-PAGE (day 5-7). Cleaved protein was rocked with 5 mL of equilibrated Ni-NTA resin at 4 ⁰C for 30 min, and poured through an open column, and washed with wash buffer 1 and 2, followed by elution buffer. All other proteins were dialyzed in their respective wash buffer 1, with no tag cleavage.

HDAC6^1–1215^ and USP16^1–823^were diluted 3-fold with Buffer A (20 mM Tris pH 8, 1 mM TCEP), and then concentrated to 5 mL and loaded onto a MonoQ 5/50 column (Buffer A: 20 mM Tris pH 8, 1 mM TCEP; Buffer B: 20 mM Tris pH 8, 1 M NaCl, 1 mM TCEP). Peak fractions were analyzed by SDS-PAGE, protein fractions were pooled and then loaded onto a S200 16/60 gel filtration column for further purification (gel filtration buffer: 50 mM Tris pH 8, 150 mM NaCl, 1 mM TCEP). USP3^1–520^, USP33^c36–825^ were concentrated to 5 mL and loaded onto a S200 16/60 gel filtration column. All other dialyzed or cleaved proteins were concentrated to 5 mL and loaded onto a S75 16/60 gel filtration column. Protein peak fractions were analyzed by SDS-PAGE, pooled and concentrated. Protein concentration was measured by UV absorbance at 280 nm, and protein identity was confirmed by mass spectrometry.

### FLUORESCENCE POLARIZATION ASSAY

Experiments were performed in 384-well black polypropylene PCR plates (Axygen) in 10 µL volume. In each well, 9 µL compound solutions in buffer (20 mM HEPES pH 7.4, 150 mM NaCl, 1 mM TCEP, 0.005% Tween-20 (v/v), 1% DMSO (v/v)) were diluted, followed by the addition of 1 µL of 30 µM HDAC6^1109–1213^ or 10 µM HDAC6^1–1215^ and 500 nM N-terminally FITC-labeled RLRGG or 1 µL of 2 µM HDAC6^1109–1213^ and 500 nM N-terminally FITC-labeled LRLRGG were added to each well. The LRLRGG substrate was used for compounds with K_D_ < ∼1 µM. Following 1 min centrifugation at 1000 RPM, the plate was incubated for 10 minutes before FP measurements with a BioTek Synergy 4 (BioTek) at excitation and emission wavelengths of 485 and 528 nm, respectively. The data was processed in GraphPad Prism using Sigmoidal, 4PL, X is log(concentration) fit.

### ISOTHERMAL TITRATION CALORIMETRY

HDAC6^1109–1215^ was diluted to 10 µM and dialyzed for 24 hours at 4 ⁰C in assay buffer (PBS, 0.005% Tween-20 (v/v), 0.25% DMSO (v/v). HDAC6^1109–1215^ concentration was assessed after dialysis using UV absorbance at 280 nm. **25** was initially diluted from the solubilized DMSO stock to 0.25% DMSO (v/v) in the assay buffer (no DMSO) for a final concentration of 100 µM. The pH of the protein and compound solution were measured and assessed to be within 0.1 pH units of each other. ITC measurements were performed at 25 ⁰C on a Nano ITC (TA Instruments), with 300 µL of 10 µM HDAC6 in the sample cell, and 50 µL of 100 µM **25** in the injection syringe. A total of 24×2 µL titrations with an initial 0.5 µL injection that was omitted from the fitting analysis were delivered into a 0.167 mL sample cells at a 180 s interval. The data was analyzed using Nano Analyze software and fitted with an independent-binding site model.

### SURFACE PLASMON RESONANCE

Studies were performed using a Biacore T200 (GE Health Sciences). A SA chip was primed with 3×60 s injection with 50 mM NaOH, followed by approximately 500-6000 response units (RU) of biotinylated protein (HDAC6^1109–1215^, HDAC6^1108–1215^ R1155A_Y1184A HDAC6^1–1215^ _USP31-131, USP5171-290, USP13183-307, USP1625-185, USP201-141, USP3329-134, USP3984-194, USP491-115,_ USP51^176–305^ and BRAP^304–390^) diluted in assay buffer (20 mM HEPES pH 7.4, 150 mM NaCl, 1 mM TCEP, 0.005% Tween-20 (v/v), and 1% DMSO (v/v)) coupling to flow channels, with an empty flow channel used for reference subtraction. Following protein capture, assay buffer was flowed over the chip until a stable baseline was achieved. Compound dilutions were prepared in assay buffer and experiments were performed using multicycle kinetics with 60-400 s contact time, 60-400s dissociation time and 20-30 µL/min flow rate at 20 ⁰C. K_D_ values were calculated using steady-state affinity fitting with the Biacore T200 evaluation software.

### X-RAY CRYSTALLOGRAPHY

Apo 3.5 mg/mL HDAC6^1109–1213^ was crystallized in mother liquor: 2 M Na formate, 0.1 M Na acetate pH 4.6, 5% ethylene glycol. 1 µL of apo HDAC6^1109–1213^ crystals were diluted 1:1000 with mother liquor and vortexed vigorously to make seed mix. Crystallization solutions of 3.5 mg/mL tag-free HDAC6^1109–1213^ and 1% (v/v) of a 200 mM DMSO-solubilized stock of 2, 5, 9 and 18 was prepared in buffer (50 mM Tris pH 8, 150 mM NaCl, 1 mM TCEP) and incubated on ice for at least 1 hour. 1 µL HDAC6^1109–1213^: compound solution and 1 µL of the mother liquor were set up in a 96-well Intelliplate using a PHOENIX liquid dispensing robot followed by the addition of 200 nL of 1:2-8-fold dilution of seed mix using a MOSQUITO instrument. X-ray diffraction data for HDAC6^1109–1213^ co-crystals were collected at 100 K at a Rigaku FR-E Superbright at a wavelength of 1.54178 Å for compound **9** and **15**. X-ray diffraction data for HDAC6^1109–1213^ co-crystal with **25** were collected at 100 K at APS 24-ID-E at a wavelength of 0.9792 Å. Diffraction data were processed with Xia2 and Aimless ^2^^6^. Models were refined with cycles of Coot ^27^ for model building and visualization, with REFMAC ^28^ for restrained refinement and validated with MOLPROBITY^29^.

### HDAC6 CATALYTIC ACTIVITY ASSAY

Final concentration of 200 nM HDAC6^1–1215^ was incubated with compound dilutions prepared with 20 mM HEPES pH 8.0, 1 mM MgCl_2_, 137 mM NaCl, 2.7 mM KCl, 0.05% BSA (v/v), 2% DMSO (v/v) in a 384-well microplate (Greiner). A final concentration of 50 µM Boc-Lys(TFA)-AMC was added and incubated for 1 hour at room temperature followed by the addition of developer solution containing a final concentration of 50 µM TSA. Following a 1-minute centrifugation at 1000 RPM, fluorescence intensity was read using BioTek Synergy 4 (BioTek) with excitation and emission of 360 and 460 nm. The data were analyzed with GraphPad Prism 8.2.0.

### UBIQUITIN RHODAMINE DUB ASSAY

Experiments were performed in a total volume of 60 μL in 384-well black polypropylene microplates (Grenier). Fluorescence was measured using a Biotek Synergy H1 microplate reader (Biotek) at excitation and emission wavelengths of 485 and 528 nm, respectively. Compound dilutions were prepared in assay buffer (USP3^1–520^: 20 mM Tris pH 7.5, 125 mM NaCl, 1 mM DTT, 0.01% TX-100 (v/v), 1% DMSO (v/v); USP5^1–835^: 20 mM Tris pH 7.5, 30 mM NaCl, 1 mM DTT, 0.01% Triton-X (v/v), and 1% DMSO (v/v); USP16^1–823^: 20 mM Tris pH 7.5, 150 mM NaCl, 1 mM DTT, 0.002% TX-100 (v/v), 1% DMSO (v/v), USP33^c36–825^: 20 mM Tris pH 7.5, 30 mM NaCl, 5 mM DTT, 0.01% TX-100 (v/v), 1% DMSO (v/v)). 500 nM USP31-520, 1 nM USP5^1−8^^c35^, 16 nM USP16^1–823^, 8 nM USP33^c36–825^) and 200 nM ubiquitin-rhodamine110 (UBPBio) (for USP3^1–520^ and USP5^1−8^^c35^), or 500 nM (for USP16^1–823^, USP33^c36–825^) was added to each well. Following 1 min centrifugation at 1000 RPM, fluorescence readings were immediately taken for 10 min. The data were analyzed with GraphPad Prism 8.2.0.

### NANOBRET ASSAY

HEK293T cells were plated in 6-well plates (8 x10^5^/well) in DMEM supplemented with 10% FBS, penicillin (100 U/mL) and streptomycin (100 µg/mL). After 4 hours cells were co-transfected with 0.02 µg N-terminally NanoLuc-tagged HDAC6 or USP16 (wildtype or R115A/R84A, Y1184A/Y117A mutant, respectively), 0.4 µg N-terminally HaloTag-tagged ISG15 and 1.6 µg of empty vector. The following day cells were trypsinized and seeded in 384-well white plate (20 µl/well) in DMEM F12 (no phenol red, 4% FBS) +/-HaloTag® NanoBRET™ 618 Ligand (1 µl/ml, Promega) and +/-compounds (DMSO concentration in each sample was kept the same). 4 h later 5 µl/well of NanoBRET™ Nano-GloR Substrate (10 µl/ml in DMEM no phenol red, Promega) was added and 460 nm donor and 618 nm acceptor signals were read within 10 min of substrate addition using CLARIOstar microplate reader (Mandel). Mean corrected NanoBRET ratios (mBU) were determined by subtracting mean of 618/460 signal from cells without NanoBRET™ 618 Ligand x 1000 from mean of 618/460 signal from cells with NanoBRET™ 618 Ligand x 1000. The IC_50_ values were determined using GraphPad Prism 7 software.

### HDAC6 PULL DOWN FROM WHOLE CELL LYSATE AND WESTERN BLOT

HEK-293 cells were cultured in DMEM supplemented with 10% fetal bovine serum (FBS) and 100 U/ml penicillin and 100 μg/ml streptomycin, in 15 cm plates. To isolate whole cell lysate, media was removed from the plate and cells were washed once in 2 mL phosphate buffered saline (PBS). Cells were scraped off the plate using a cell scraper, transferred into a 15 mL tube, and centrifuged for 3 minutes at 230 g. Cells were lysed in 1 mL lysis buffer with salt (450 mM NaCl, 50 mM Tris-HCl pH 7.5, 1 mM EDTA, 1% Triton X-100, 1X protease inhibitors) and mixed by vigorous pipetting and vortexing. Cells were incubated on ice for 20 minutes before dilution with lysis buffer without salt (50 mM Tris-HCl pH 7.5, 1 mM EDTA, 1% Triton X-100, 1X protease inhibitors) to a final concentration of 150 mM NaCl. Lysates were centrifuged at 20,000 g for 2 minutes at 4 ⁰C and supernatant was transferred to a fresh tube. Protein concentration was determined using Pierce BCA Protein Assay and lysates were snap frozen in liquid nitrogen and stored at -80 ⁰C. Lysates were thawed and a small volume was set aside as an input control and stored at -20⁰C. To each 3 mg of thawed lysate, **25** or **32** or DMSO control (same volume of DMSO) was added to achieve a final compound concentration of 10 µM. Tubes were incubated with rotation for 1 hour at 4 ⁰C. 25 µL Streptavidin magnetic Dynabeads (M270, Thermo-Fischer Scientific 65305) per sample were combined into a single tube and washed twice in 500 µL low salt wash buffer (100 mM NaCl, 10 mM Tris-HCl pH 7.9, 0.1% NP-40), with beads isolated from the buffer between each step using a magnetic rack. An aliquot of 25 µL of washed beads was removed and added to lysate without pre-incubation with any compounds or DMSO (beads alone control). The remaining washed beads were incubated for 1 hour at 4 ⁰C with rotation in 2 mL low salt wash buffer containing 10 µM **33**. Excess unbound compound was removed by washing the beads three times in 1 mL low salt wash buffer before resuspending the beads in 100 µL low salt wash buffer per 25 µL of beads. After 1 hour pre-incubation of lysates with DMSO, **25**, or **32**, 100 µL **33**-bound beads were added and lysates were incubated for 1 hour with rotation at 4 ⁰C. Beads were washed three times in low salt wash buffer. After the final wash, beads were pelleted and resuspended directly in 15 µl 1 x SDS loading buffer and boiled at 95 ⁰C for 1 minute. Beads were pelleted and the entire volume of supernatant was used for western blotting along with input lysate using the NuPAGE electrophoresis and transfer system (Invitrogen) and near-infrared detection for HDAC6 (CST #7558; 1:1000) using IRDye® 800CW Secondary Antibody (1:5000). Immunoblots were imaged on a Li-Cor Odyssey CLx.

### HDAC6 CHEMOPROTEOMICS IN CYTOPLASMIC FRACTION

To isolate the cytoplasmic fraction, cell pellets were collected as above and resuspended in 3 mL hypotonic lysis buffer, Buffer A (10 mM HEPES pH 7.4, 10 mM NaCl, 1.5 mM MgCl_2_, 0.005% (v/v) Tween-20, 1X protease inhibitors), and kept on ice for 20 minutes, with vigorous vortexing for 5 seconds every 5 minutes. Cells were centrifuged for 5 minutes at 380 g and 4 ⁰C, supernatant was collected and transferred to 2 mL tubes before being centrifuged again for 1 minute at 18,500 g and 4 ⁰C. Cleared supernatant was collected and transferred to 2 mL tubes. NaCl was added to adjust the final concentration to 150 mM. Protein concentration was determined using Pierce BCA Protein Assay and lysates were snap frozen in liquid nitrogen and stored at -80 ⁰C.

25 nanomoles of **33** was bound to 20 µL of Streptavidin Sepharose® High Performance (MilliporeSigma, GE17-5113-01) beads for 1 h at 4 ⁰C in PBS. The beads were washed 3x with Buffer A. Meanwhile, to each 1.5 mg of cytoplasmic fraction, a final compound concentration of 10 µM compound 25, 32 or DMSO control (same volume of DMSO) was added and samples incubated for 1 h at 4 ⁰C. Protein and beads were then mixed and rocked for a further 1h at 4 ⁰C. The supernatant was removed and the beads were washed 1 time with Buffer A and transferred to a new tube. The beads were then washed beads 3x with 50 mM ammonium bicarbonate and then 1 µg chymotrypsin was added for 15 min at RT. To this solution, 1 µg trypsin was added and incubated for 2.25 h at 37 ⁰C. Disulfide bonds were reduced by adding DTT to a final concentration of 5 mM. After incubating for 30 minutes at 56 ⁰C, the reduced cysteines were alkylated with 20 mM iodoacetamide in the dark for 45 min. An additional 1 µg trypsin was added and the solution was left overnight at 37 ⁰C.

The digested peptides were analyzed using reversed-phase (Reprosil-Pur 120 C18-AQ, 1.9 µm), nano-HPLC (Vanquish Neo UHPLC) coupled to an Orbitrap Fusion™ Lumos™ Tribrid™. Peptides were eluted from the column with an acetonitrile gradient starting from 3.2% acetonitrile with 0.1% formic acid to 35.2% acetonitrile with 0.1% formic acid using a linear gradient of 90 minutes. The MS1 scan had an accumulation time of 50 ms within a mass range of 400–1500Da, using orbitrap resolution of 120000, 60% RF lens, AGC target of 125% and 2400 volts. This was followed by MS/MS scans with a total cycle time of 3 seconds. Accumulation time of 50 ms and 33% HCD collision energy was used for each MS/MS scan. Each candidate ion was required to have a charge state from 2-7 and an AGC target of 400%, isolated using orbitrap resolution of 15,000. Previously analyzed candidate ions were dynamically excluded for 9 seconds. The RAW files were searched with FragPipe v18.0, using MSFragger v3.5 and Philosopher v4.4.0 ^30, 31^. Utilized the LFQ-MBR workflow using chymotrypsin/trypsin enzymatic digestion with human Uniprot ID UP000005640 (with decoys and contaminants appended). Differential protein expression was determined using R Package DEP ^24^, and independently validated with ProDA ^25^.

## ASSOCIATED CONTENT

### Supporting Information

Additional data regarding supplementary tables and figures. Data collection and refinement statistics for compounds **9**, **15** and **25** (**Table S2**). General practical considerations, synthetic procedures, characterization data and purity analysis for compounds investigated in this study.

Molecular formula strings and the associated biological data Crystallographic information files

### Accession Codes

PDB deposition: HDAC6 UBD in complex with **9** – 8G43, HDAC6 UBD in complex with **15** – 8G44, HDAC6 UBD in complex with **25** – 8G45. Authors will release the atomic coordinates and experimental data upon article publication.

Mass spectrometry data deposition: Data are available via ProteomeXchange with identifier PXD039880. Processed mass spectrometry data files and other associated materials are available via Zenodo #7618095. Files in the deposition are as follows:

| Filename | Description |
| --- | --- |
| combined_protein_Int.txt | Output from database search |
| hdac6_dep_submitted.R | script used to process data (from DEP package) |
| hdac6_dep_submitted.R.RData | workspace environment from DEP package |
| hdac6probecontrol_dep.csv | exported list of p-value and fold-change from DEP |
| hdac6probecontrol_proda.csv | exported list of p-value and fold-change from proDA |
| hdac6proda_submitted.R | proDA script |
| hdac6proda_submitted.RData | proDA workspace environment |

## AUTHOR INFORMATION

### Corresponding Authors

Mark Lautens

Cheryl H. Arrowsmith

### Author Contributions

R.J.H., I.F, M.K.M, M.S. and B.M contributed equally to this work. V.S, M.S designed compounds, R.J.H wrote the manuscript, R.J.H solved structures, R.J.H, M.K.M, M.S, S.A and D.O tested compounds, and I.F, B.M, A.S., K.J., R.S., R.B and C.D synthesized compounds. C.H.A. conceived the project and C.H.A., M.L and P.J.B. advised throughout the project. All authors have given approval to the final version of the manuscript.

## ACKNOWLEDGEMENTS

The Structural Genomics Consortium is a registered charity (no: 1097737) that receives funds from Bayer AG, Boehringer Ingelheim, Bristol Myers Squibb, Genentech, Genome Canada through Ontario Genomics Institute [OGI-196], EU/EFPIA/OICR/McGill/KTH/Diamond Innovative Medicines Initiative 2 Joint Undertaking [EUbOPEN grant 875510], Janssen, Merck KGaA (aka EMD in Canada and US), Pfizer and Takeda. I.F, B.M, C.D, R.S were supported by NSERC CREATE grant. C.D was supported by an Erwin Schroedinger postdoctoral fellowship, awarded by the Austrian Science Fund (FWF): J 4348-N28. The authors acknowledge Dr Matthew Maitland for constructive discussions about chemoproteomics. Proteomics experiments were done at the Network Biology Collaborative Centre at the Lunenfeld-Tanenbaum Research Institute. This work is based upon research conducted at the Northeastern Collaborative Access Team beamlines, which are funded by the National Institute of General Medical Sciences from the National Institutes of Health (P30 GM124165). The Eiger 16M detector on the 24-ID-E beamline is funded by a NIH-ORIP HEI grant (S10OD021527). This research used resources of the Advanced Photon Source, a U.S. Department of Energy (DOE) Office of Science User Facility operated for the DOE Office of Science by Argonne National Laboratory under Contract No. DE-AC02-06CH11357.

## ABBREVIATIONS

HDAC: Histone deacetylase
UBD: ubiquitin binding domain
K_D_: dissociation constant
SPR: surface plasmon resonance
FP: fluorescence polarisation
NanoBRET: nano-bioluminescence resonance energy transfer

## REFERENCES

(1) de Ruijter, A. J. M.; van Gennip, A. H.; Caron, H. N.; Kemp, S.; van Kuilenburg, A. B. P. Histone Deacetylases (HDACs): Characterization of the Classical HDAC Family. The Biochemical journal 2003, 370 (Pt 3), 737–749. https://doi.org/10.1042/BJ20021321.

(2) Hubbert, C.; Guardiola, A.; Shao, R.; Kawaguchi, Y.; Ito, A.; Nixon, A.; Yoshida, M.; Wang, X.-F.; Yao, T.-P. HDAC6 Is a Microtubule-Associated Deacetylase. Nature 2002, 417 (6887), 455–458. https://doi.org/10.1038/417455a.

(3) Hai, Y.; Christianson, D. W. Histone Deacetylase 6 Structure and Molecular Basis of Catalysis and Inhibition. Nature chemical biology 2016, 12 (9), 741–747. https://doi.org/10.1038/nchembio.2134.

(4) Hook, S. S.; Orian, A.; Cowley, S. M.; Eisenman, R. N. Histone Deacetylase 6 Binds Polyubiquitin through Its Zinc Finger (PAZ Domain) and Copurifies with Deubiquitinating Enzymes. Proceedings of the National Academy of Sciences of the United States of America 2002. https://doi.org/10.1073/pnas.172511699.

(5) Boyault, C.; Gilquin, B.; Zhang, Y.; Rybin, V.; Garman, E.; Meyer-Klaucke, W.; Matthias, P.; Müller, C. W.; Khochbin, S. HDAC6-P97/VCP Controlled Polyubiquitin Chain Turnover. The EMBO journal 2006, 25 (14), 3357–3366. https://doi.org/10.1038/sj.emboj.7601210.

(6) Kawaguchi, Y.; Kovacs, J. J.; McLaurin, A.; Vance, J. M.; Ito, A.; Yao, T.-P. The Deacetylase HDAC6 Regulates Aggresome Formation and Cell Viability in Response to Misfolded Protein Stress. Cell 2003, 115 (6), 727–738. https://doi.org/10.1016/S0092-8674(03)00939-5.

(7) Hideshima, T.; Bradner, J. E.; Wong, J.; Chauhan, D.; Richardson, P.; Schreiber, S. L.; Anderson, K. C. Small-Molecule Inhibition of Proteasome and Aggresome Function Induces Synergistic Antitumor Activity in Multiple Myeloma. Proc. Natl. Acad. Sci. U.S.A. 2005, 102 (24), 8567–8572. https://doi.org/10.1073/pnas.0503221102.

(8) Miyake, Y.; Keusch, J. J.; Wang, L.; Saito, M.; Hess, D.; Wang, X.; Melancon, B. J.; Helquist, P.; Gut, H.; Matthias, P. Structural Insights into HDAC6 Tubulin Deacetylation and Its Selective Inhibition. Nat. Chem. Biol. 2016, 12 (9), 748–754. https://doi.org/10.1038/nchembio.2140.

(9) Nakashima, H.; Nguyen, T.; Goins, W. F.; Chiocca, E. A. Interferon-Stimulated Gene 15 (ISG15) and ISG15-Linked Proteins Can Associate with Members of the Selective Autophagic Process, Histone Deacetylase 6 (HDAC6) and SQSTM1/P62. J Biol Chem 2015, 290 (3), 1485–1495. https://doi.org/10.1074/jbc.M114.593871.

(10) Ouyang, H.; Ali, Y. O.; Ravichandran, M.; Dong, A.; Qiu, W.; MacKenzie, F.; Dhe-Paganon, S.; Arrowsmith, C. H.; Zhai, R. G. Protein Aggregates Are Recruited to Aggresome by Histone Deacetylase 6 via Unanchored Ubiquitin C Termini. J. Biol. Chem. 2012, 287 (4), 2317–2327. https://doi.org/10.1074/jbc.M111.273730.

(11) Brindisi, M.; Saraswati, A. P.; Brogi, S.; Gemma, S.; Butini, S.; Campiani, G. Old but Gold: Tracking the New Guise of Histone Deacetylase 6 (HDAC6) Enzyme as a Biomarker and Therapeutic Target in Rare Diseases. J. Med. Chem. 2020, 63 (1), 23–39. https://doi.org/10.1021/acs.jmedchem.9b00924.

(12) Simões-Pires, C.; Zwick, V.; Nurisso, A.; Schenker, E.; Carrupt, P.-A.; Cuendet, M. HDAC6 as a Target for Neurodegenerative Diseases: What Makes It Different from the Other HDACs? Molecular Neurodegeneration 2013, 8 (1), 7. https://doi.org/10.1186/1750-1326-8-7.

(13) Li, T.; Zhang, C.; Hassan, S.; Liu, X.; Song, F.; Chen, K.; Zhang, W.; Yang, J. Histone Deacetylase 6 in Cancer. Journal of Hematology & Oncology 2018, 11 (1), 111. https://doi.org/10.1186/s13045-018-0654-9.

(14) Zeleke, T. Z.; Pan, Q.; Chiuzan, C.; Onishi, M.; Li, Y.; Tan, H.; Alvarez, M. J.; Honan, E.; Yang, M.; Chia, P. L.; Mukhopadhyay, P.; Kelly, S.; Wu, R.; Fenn, K.; Trivedi, M. S.; Accordino, M.; Crew, K. D.; Hershman, D. L.; Maurer, M.; Jones, S.; High, A.; Peng, J.; Califano, A.; Kalinsky, K.; Yu, J.; Silva, J. Network-Based Assessment of HDAC6 Activity Predicts Preclinical and Clinical Responses to the HDAC6 Inhibitor Ricolinostat in Breast Cancer. Nat Cancer 2022, 1–19. https://doi.org/10.1038/s43018-022-00489-5.

(15) Lee, E. K.; Tan-Wasielewski, Z.; Matulonis, U. A.; Birrer, M. J.; Wright, A. A.; Horowitz, N.; Konstantinopoulos, P. A.; Curtis, J.; Liu, J. F. Results of an Abbreviated Phase Ib Study of the HDAC6 Inhibitor Ricolinostat and Paclitaxel in Recurrent Ovarian, Fallopian Tube, or Primary Peritoneal Cancer. Gynecologic Oncology Reports 2019, 29, 118–122. https://doi.org/10.1016/j.gore.2019.07.010.

(16) Vogl, D. T.; Raje, N.; Jagannath, S.; Richardson, P.; Hari, P.; Orlowski, R.; Supko, J. G.; Tamang, D.; Yang, M.; Jones, S. S.; Wheeler, C.; Markelewicz, R. J.; Lonial, S. Ricolinostat, the First Selective Histone Deacetylase 6 Inhibitor, in Combination with Bortezomib and Dexamethasone for Relapsed or Refractory Multiple Myeloma. Clin. Cancer Res. 2017. https://doi.org/10.1158/1078-0432.CCR-16-2526.

(17) Yee, A. J.; Bensinger, W. I.; Supko, J. G.; Voorhees, P. M.; Berdeja, J. G.; Richardson, P. G.; Libby, E. N.; Wallace, E. E.; Birrer, N. E.; Burke, J. N.; Tamang, D. L.; Yang, M.; Jones, S. S.; Wheeler, C. A.; Markelewicz, R. J.; Raje, N. S. Ricolinostat plus Lenalidomide, and Dexamethasone in Relapsed or Refractory Multiple Myeloma: A Multicentre Phase 1b Trial. The Lancet Oncology 2016, 17 (11), 1569–1578. https://doi.org/10.1016/S1470-2045(16)30375-8.

(18) Suzuki, T.; Miyata, N. Non-Hydroxamate Histone Deacetylase Inhibitors. Curr. Med. Chem. 2005, 12 (24), 2867–2880.

(19) Harding, R. J.; Ferreira De Freitas, R.; Collins, P.; Franzoni, I.; Ravichandran, M.; Ouyang, H.; Juarez-Ornelas, K. A.; Lautens, M.; Schapira, M.; Von Delft, F.; Santhakumar, V.; Arrowsmith, C. H. Small Molecule Antagonists of the Interaction between the Histone Deacetylase 6 Zinc-Finger Domain and Ubiquitin. Journal of Medicinal Chemistry 2017. https://doi.org/10.1021/acs.jmedchem.7b00933.

(20) Ferreira De Freitas, R.; Harding, R. J.; Franzoni, I.; Ravichandran, M.; Mann, M. K.; Ouyang, H.; Lautens, M.; Santhakumar, V.; Arrowsmith, C. H.; Schapira, M. Identification and Structure-Activity Relationship of HDAC6 Zinc-Finger Ubiquitin Binding Domain Inhibitors. Journal of Medicinal Chemistry 2018. https://doi.org/10.1021/acs.jmedchem.8b00258.

(21) Robers, M. B.; Vasta, J. D.; Corona, C. R.; Ohana, R. F.; Hurst, R.; Jhala, M. A.; Comess, K. M.; Wood, K. V. Quantitative, Real-Time Measurements of Intracellular Target Engagement Using Energy Transfer. Methods Mol Biol 2019, 1888, 45–71. https://doi.org/10.1007/978-1-4939-8891-4_3.

(22) Dao, C. T.; Zhang, D.-E. ISG15: A Ubiquitin-like Enigma. Front Biosci 2005, 10, 2701– 2722. https://doi.org/10.2741/1730.

(23) Sen Nkwe, N.; Daou, S.; Uriarte, M.; Gagnon, J.; Iannantuono, N. V.; Barbour, H.; Yu, H.; Masclef, L.; Fernández, E.; Zamorano Cuervo, N.; Mashtalir, N.; Binan, L.; Sergeev, M.; Bélanger, F.; Drobetsky, E.; Milot, E.; Wurtele, H.; Costantino, S.; Affar, E. B. A Potent Nuclear Export Mechanism Imposes USP16 Cytoplasmic Localization during Interphase. J Cell Sci 2020, 133 (4), jcs239236. https://doi.org/10.1242/jcs.239236.

(24) Zhang, X.; Smits, A. H.; van Tilburg, G. B.; Ovaa, H.; Huber, W.; Vermeulen, M. Proteome-Wide Identification of Ubiquitin Interactions Using UbIA-MS. Nat Protoc 2018, 13 (3), 530– 550. https://doi.org/10.1038/nprot.2017.147.

(25) Ahlmann-Eltze, C.; Anders, S. ProDA: Probabilistic Dropout Analysis for Identifying Differentially Abundant Proteins in Label-Free Mass Spectrometry. bioRxiv May 1, 2020, p 661496. https://doi.org/10.1101/661496.

(26) Evans, P. R.; Murshudov, G. N. How Good Are My Data and What Is the Resolution? Acta Crystallogr D Biol Crystallogr 2013, 69 (Pt 7), 1204–1214. https://doi.org/10.1107/S0907444913000061.

(27) Emsley, P.; Lohkamp, B.; Scott, W. G.; Cowtan, K. Features and Development of Coot. Acta Crystallogr D Biol Crystallogr 2010, 66 (Pt 4), 486–501. https://doi.org/10.1107/S0907444910007493.

(28) Murshudov, G. N.; Skubák, P.; Lebedev, A. A.; Pannu, N. S.; Steiner, R. A.; Nicholls, R. A.; Winn, M. D.; Long, F.; Vagin, A. A. REFMAC5 for the Refinement of Macromolecular Crystal Structures. Acta Cryst D 2011, 67 (4), 355–367. https://doi.org/10.1107/S0907444911001314.

(29) Williams, C. J.; Headd, J. J.; Moriarty, N. W.; Prisant, M. G.; Videau, L. L.; Deis, L. N.; Verma, V.; Keedy, D. A.; Hintze, B. J.; Chen, V. B.; Jain, S.; Lewis, S. M.; Arendall, W. B.; Snoeyink, J.; Adams, P. D.; Lovell, S. C.; Richardson, J. S.; Richardson, D. C. MolProbity: More and Better Reference Data for Improved All-atom Structure Validation. Protein Sci 2018, 27 (1), 293–315. https://doi.org/10.1002/pro.3330.

(30) da Veiga Leprevost, F.; Haynes, S. E.; Avtonomov, D. M.; Chang, H.-Y.; Shanmugam, A. K.; Mellacheruvu, D.; Kong, A. T.; Nesvizhskii, A. I. Philosopher: A Versatile Toolkit for Shotgun Proteomics Data Analysis. Nat Methods 2020, 17 (9), 869–870. https://doi.org/10.1038/s41592-020-0912-y.

(31) Kong, A. T.; Leprevost, F. V.; Avtonomov, D. M.; Mellacheruvu, D.; Nesvizhskii, A. I. MSFragger: Ultrafast and Comprehensive Peptide Identification in Mass Spectrometry-Based Proteomics. Nat Methods 2017, 14 (5), 513–520. https://doi.org/10.1038/nmeth.4256.

