## Supplementary Information for "Discovery and characterization of a chemical probe targeting the zinc-finger ubiquitin-binding domain of HDAC6"

SUPPORTING INFORMATION

**Table S1.** Affinity of HDAC6 proteins for ubiquitin and ISG substrates used in this study

| **Substrate** | **HDAC6 ZnF-UBD** | | **HDAC6 ZnF-UBD R1155A Y1184A** | **Full-Length HDAC6** |
| --- | --- | --- | --- | --- |
|  | **FP K_D_ (µM)*^a^*** | **SPR K_D_ (µM)***^b^* | **SPR K_D_ (µM) *^b^*** | **SPR K_D_ (µM) *^b^*** |
| **RLRGG peptide** | NT | 11 ± 4 | NT | 40 ± 25 |
| **LRLRGG peptide** | NT | 4 ± 0.7 | NT | 10 ± 7 |
| **FITC-RLRGG peptide** | 0.22 ± 0.01 | 0.33 ± 0.05 | NB | 0.23 ± 0.06 |
| **FITC-LRLRGG peptide** | 0.04 ± 0.02 | 0.33 ± 0.15 | NB | 0.8 ± 0.3 |
| **Ubiquitin** | NT | 3.2 ± 0.2 | NB | 1.0 ± 0.1 |
| **ISG15** | NT | 9.3 ± 0.6 | NB | 11 ± 2.3 |

^a^K_D_ determination experiments were performed with N=2 and the values are presented as mean ± SD. ^b^K_D_ determination experiments were performed with N=3 and the values are presented as mean ± SD. NT= not tested. NB= no binding.

**Table S2**. Data collection and refinement statistics for HDAC6 co-crystal structures.

| **Compound** | **9** | **15** | **25** |
| --- | --- | --- | --- |
| **PDB ID** | **8G43** | **8G44** | **8G45** |
| Space group | P2_1_2_1_2_1_ | P2_1_2_1_2_1_ | P2_1_2_1_2_1_ |
| a,b,c [Å] | 40.87, 44.04, 55.87 | 40.95, 44.43, 56.13 | 40.61, 44.91, 55.70 |
| α,β,γ [°] | 90.00, 90.00, 90.00 | 90.00, 90.00, 90.00 | 90.00, 90.00, 90.00 |
| Resolution limits [Å] | 29.98-1.55 (1.58-1.55) | 33.08-1.55 (1.58-1.55) | 55.70-1.62 (1.65-1.62) |
| Rmerge | 0.042 (0.254) | 0.041 (0.216) | 0.053 (0.551) |
| I/sigma | 30.7 (7.9) | 28.2 (7.9) | 14.6 (3.4) |
| Rpim | 0.018 (0.112) | 0.025 (0.139) | 0.030 (0.317) |
| CC1/2 | 1.000 (0.974) | 0.999 (0.964) | 0.999 (0.923) |
| Completeness [%] | 99.9 (100.0) | 97.9 (94.6) | 99.7 (99.2) |
| Multiplicity | 6.6 (6.1) | 6.7 (6.2) | 7.1 (7.5) |
| No. Reflections used/free | 15157/760 | 14708 (712) | 12739/681 |
| Rwork/Rfree | 0.161/0.177 | 0.166/0.198 | 0.160/0.190 |
| No. Atoms/B-factors [Å^2^] |  |  |  |
| Protein | 787/12.4 | 771/9.7 | 771/31.1 |
| Ligand | 21/14.0 | 27/8.6 | 30/34.8 |
| Water | 89/22.3 | 106/19.0 | 56/39.6 |
| Rmsd bond angle [°]/Rmsd bond length [Å] | 1.850/0.028 | 1.975/0.016 | 1.689/0.011 |
| Avg B-factors [Å^2^] | 13 | 11 | 31 |
| Molprobity Ramachandran favored/outliers [%] | 97/2 | 93/1 | 95/1 |

Values for outer shell in brackets. Omit maps for each ligand are shown in **Figure S1**.

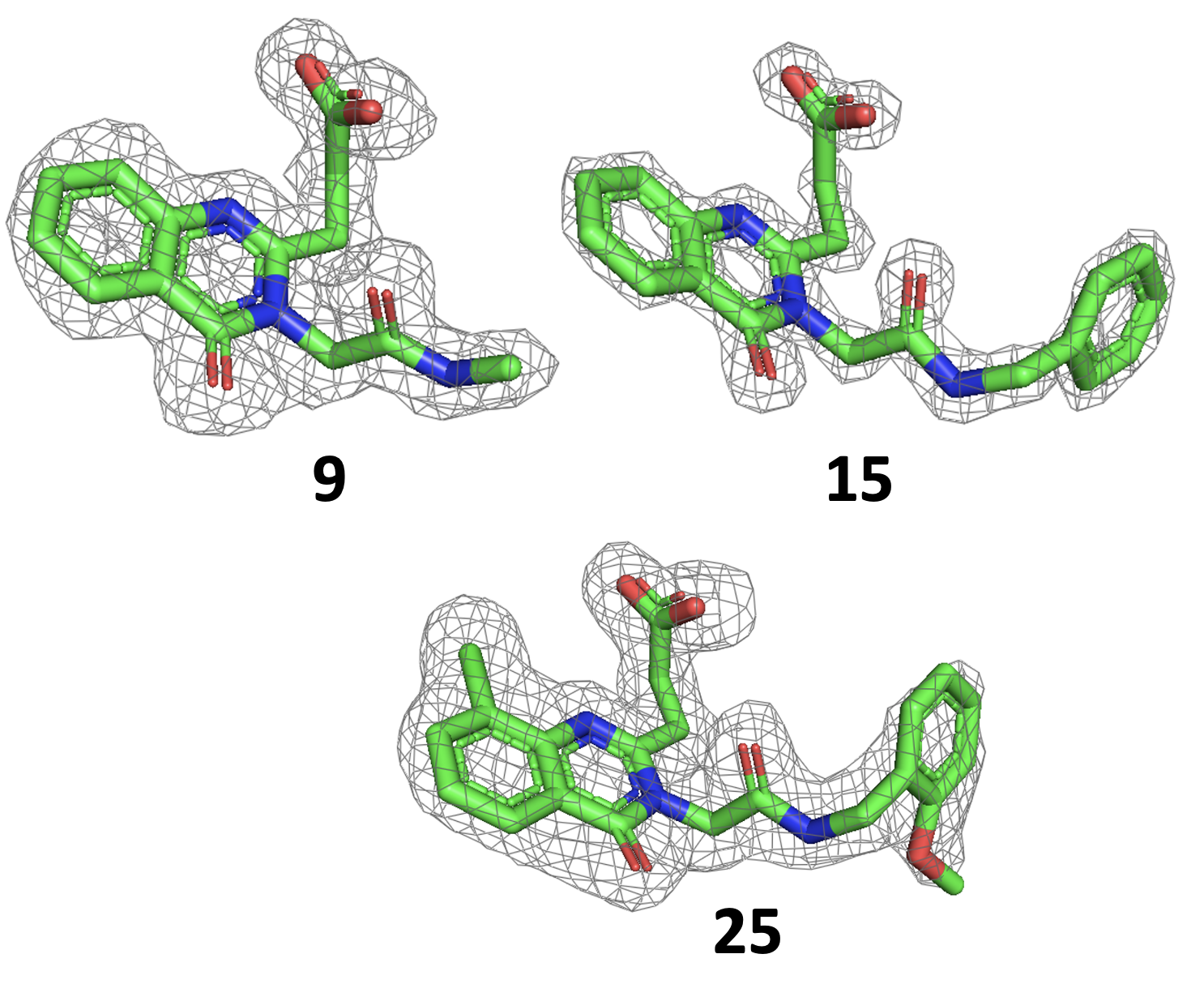

**Figure S1**. Omit map (σ2) of compounds from HDAC6-UBD crystal structures determined during structure-based optimization.

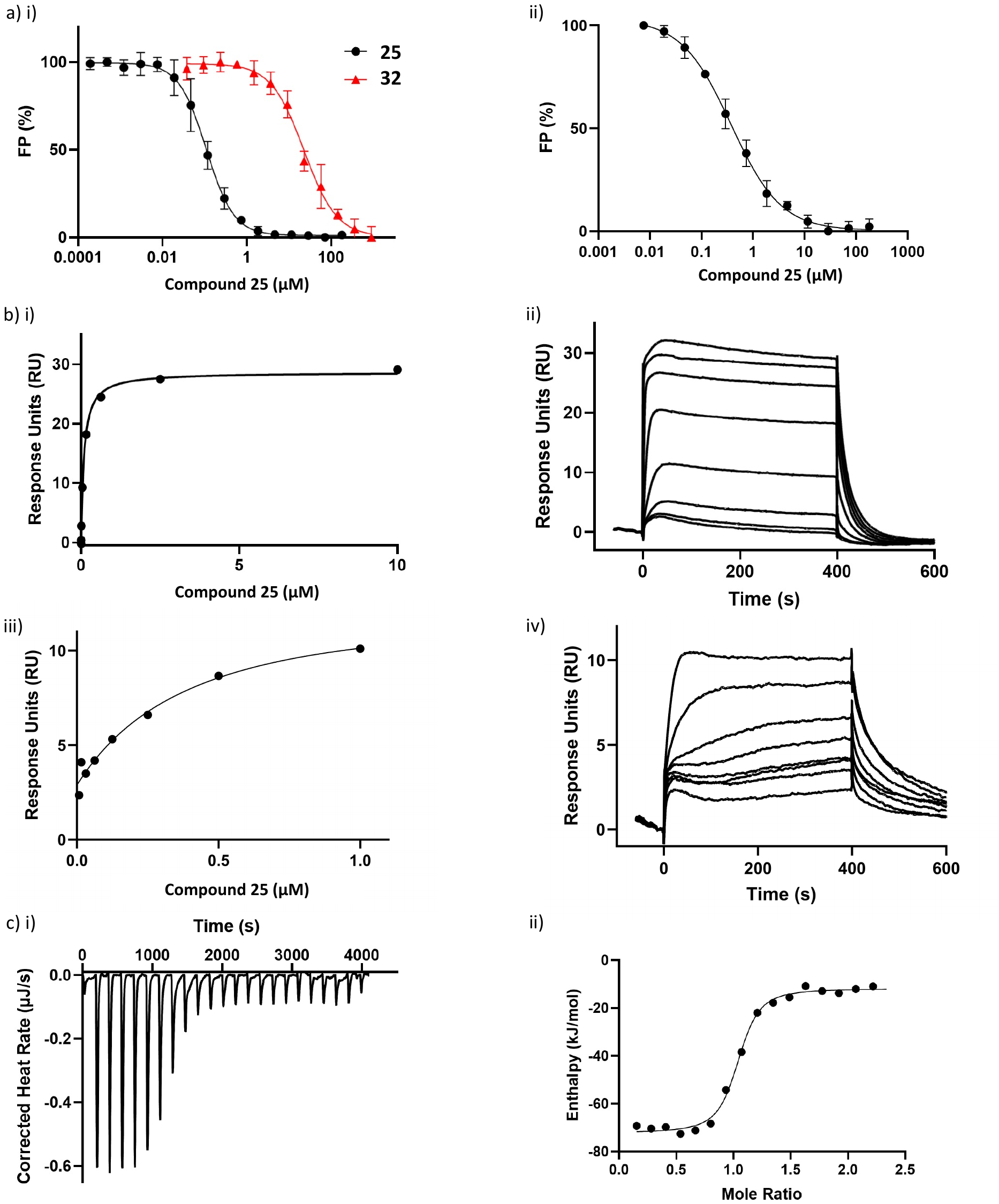

**Figure S2.** Biophysical characterization of **25** and **32**. a)i) Representative FP competition assay using increasing concentrations of **25** or **32**, a FITC-labeled LRLRGG peptide (50 nM) and HDAC6-UBD. A K_disp_ of 0.095 ± 0.018 µM, and 33 ± 6 µM was obtained from the average of three independent measurements for **25** and **32**, respectively.

**Figure S2 continued.** ii) Representative FP competition assay using increasing concentrations of **25**, a FITC-labeled LRLRGG peptide (50 nM) and full-length HDAC6. A K_disp_ of 0.26 ± 0.15 µM was obtained from the average of three independent measurements. b) Representative SPR binding i) curve and ii) sensorgram for HDAC6-UBD and **25**. A K_D_ of 0.084 ± 0.020 µM was obtained from the average of 22 independent measurements. Representative SPR binding iii) curve and iv) sensorgram for full-length HDAC6 and **25**. A K_D_ of 0.44 ± 0.09 µM was obtained from the average of 11 independent measurements. c) Representative ITC titration and fitted data showing the binding of **25** (100 µM) to HDAC6-UBD (10 µM). A K_D_ of 0.080 ± 0.023 µM was obtained from the average of three independent measurements.

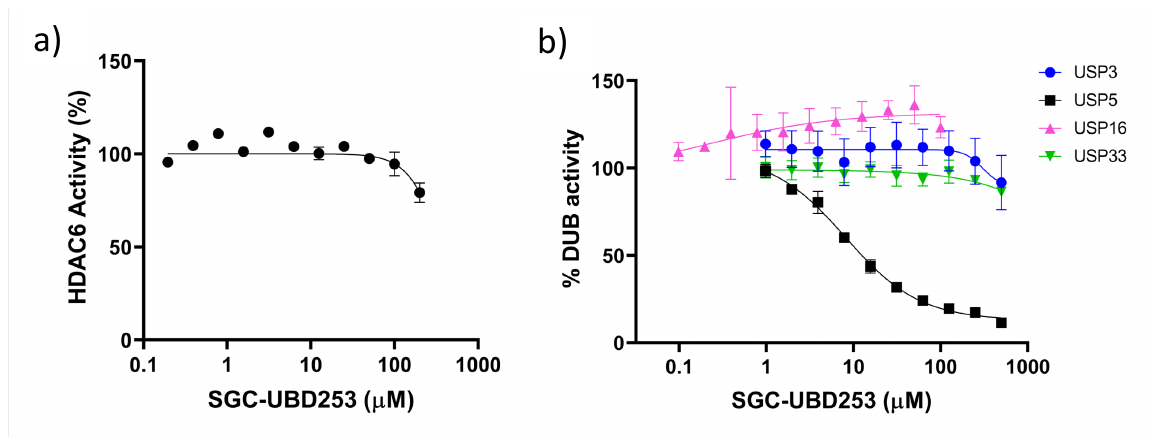

**Figure S3.** Inhibitory activity of **25**. a) HDAC6 catalytic activity assay shows **25** has no significant effect at concentrations > 100 µM (N=2) b) Ubiquitin rhodamine catalytic activity assay shows **25** does not inhibit USP3, USP16, and USP33 deubiquitinase activity but is inhibitory for USP5 activity (IC_50_= 8 ± 1 µM) (N=3).

**Synthesis of the biotinylated compound (33)**

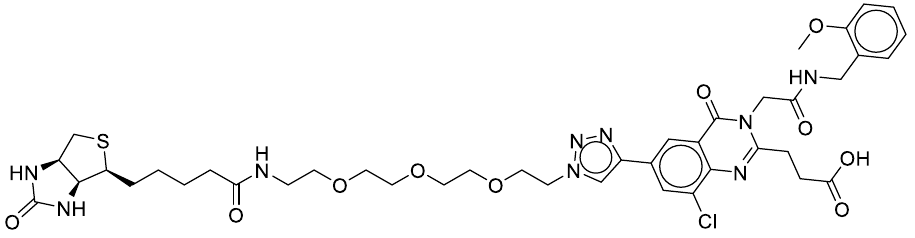

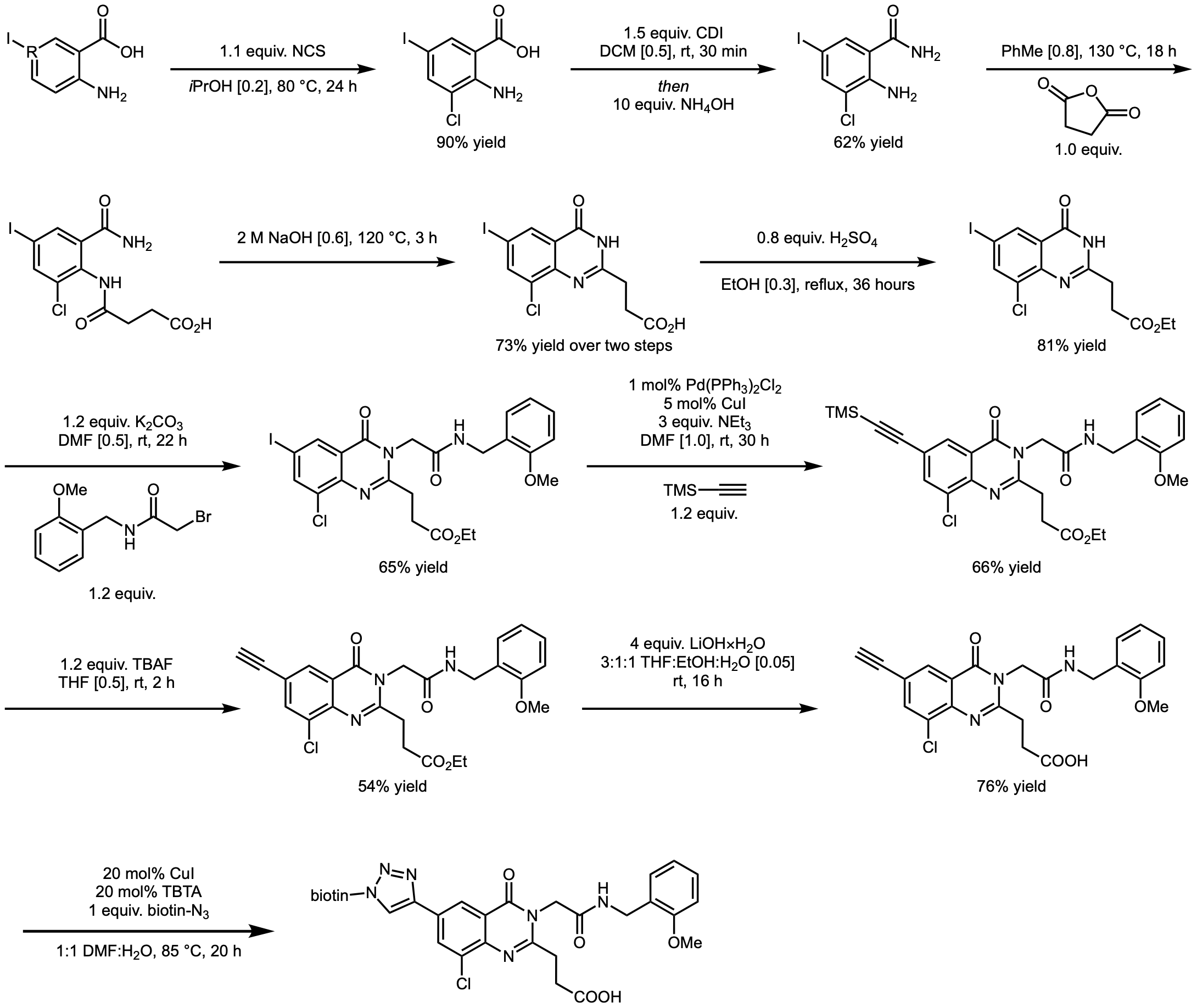

This reaction did not use glassware that was flame- or oven-dried. The iodoanthranilic acid (5.4658 g, 20.78 mmol, 1 equiv.) and NCS (3.0517 g, 22.85 mmol, 1.1 equiv.) were added to a round-bottomed flask equipped with a stir bar. Isopropanol (104 mL) was added and the solution was stirred at 80 °C for 24 hours. The solution was concentrated under reduced pressure and EtOAc was added. The organic layer was washed once with saturated sodium bisulphate, once with H_2_O, and once with brine. The organic layer was then dried over MgSO_4_ and filtered; the filtrate was concentrated *in vacuo* to afford the product as a brown solid (5.5634 g, 90% yield). The material was used as is in the next step.

The chloroiodoanthranilic acid (1.0000 g, 3.36 mmol, 1 equiv.) and CDI (817.7 mg, 5.04 mmol, 1.5 equiv.) were added to a round-bottomed flask containing a magnetic stir bar that was sealed with a septum containing an argon balloon. DCM (6.7 mL) was added and the solution was stirred at room temperature for 30 minutes. NH_4_OH (1.31 mL, 33.6 mmol, 10 equiv.) was added dropwise at room temperature and the solution was stirred for 16 hours. The solution was concentrated under reduced pressure and the residue was dissolved in EtOAc and H_2_O was added. The organics were washed twice with 1 M HCl, once with saturated NaHCO_3_, and once with brine. The organic layer was dried over MgSO_4_ and filtered; the filtrate was concentrated *in vacuo*. The crude material was recrystallized from EtOH/H_2_O to afford the chloroiodoanthranilamide as an orange solid (621.1 mg, 62% yield).

The chloroiodoanthranilamide (1.5142 g, 5.11 mmol, 1 equiv.) and succinic anhydride (511.1 mg, 5.11 mmol, 1 equiv.) were added to a round-bottomed flask containing a magnetic stir bar. Toluene (6.4 mL) was added and the mixture was stirred at 130 °C for 18 hours. The mixture was cooled to room temperature and the solids were filtered by suction and washed five times with Et_2_O. The white precipitate was collected and used as is in the next step.

This reaction did not use glassware that was flame- or oven-dried. The material from the previous step was added to a round-bottomed flask containing a magnetic stir bar and dissolved in 2 M NaOH (7 mL). The solution was stirred at 120 °C for 3 hours. The solution was then cooled to room temperature and concentrated HCl was added dropwise with stirring until the solution was acidic (pH paper). The off-white solids were collected by filtration and washed three times with H_2_O to afford the acid (1.4008 g, 73% yield over two steps).

This reaction did not use glassware that was flame- or oven-dried. The quinazolinone (14.1980 g, 37.57 mmol, 1 equiv.) was added to a round-bottomed flask containing a magnetic stir bar and suspended in EtOH (125 mL). 1.63 mL of concentrated H_2_SO_4_ was added and the mixture was refluxed for 36 hours. The solution was cooled to 0 °C, filtered, and then washed with H_2_O three times to afford the corresponding ester as a white solid (12.3598 g, 81% yield).

To a round-bottomed flask containing a magnetic stir bar was sequentially added the ester (1.0000 g, 2.46 mmol, 1 equiv.), K_2_CO_3_ (407.9 mg, 2.95 mmol, 1.2 equiv.), and the α-bromoamide (761.8 mg, 2.95 mmol, 1.2 equiv.), followed by DMF (4.9 mL). The solution was stirred at room temperature for 22 hours. The reaction was diluted with EtOAc, washed once with water and seven times with brine. The organic layer was dried over MgSO_4_ and filtered; the filtrate was concentrated *in vacuo* to afford the product as a yellow solid (932.0 mg, 65% yield) which was used as is in the next step.

This reaction did not use glassware that was flame- or oven-dried. The material from the previous step (805.2 mg, 1.38 mmol, 1 equiv.), Pd(PPh_3_)_2_Cl_2_ (9.7 mg, 0.014 mmol, 1 mol%), and CuI (13.1 mg, 0.069 mmol, 5 mol%) were added to a 2-dram vial. DMF (1.4 mL) then NEt_3_ (0.58 mL) were added sequentially, followed by TMS-acetylene (0.23 mL, 1.66 mmol, 1.2 equiv.) and the solution was stirred at room temperature for 30 hours. The mixture was filtered through a short plug of silica gel, eluting with EtOAc. The filtrate was washed twice with H_2_O, three times with brine, and dried over MgSO_4_. The mixture was filtered through a second plug of silica gel, eluting with EtOAc. The material was purified via flash column chromatography (40% EtOAc/pent, dry loaded with silica gel) to afford the product as an off-white solid (412.7 mg, 66% yield).

This reaction did not use glassware that was flame- or oven-dried. The TMS-protected alkyne (412.7 mg, 0.74 mmol, 1 equiv.) was added to a 2-dram vial and dissolved in THF (1.5 mL). A 1.0 M solution of TBAF in THF (0.89 mL, 0.89 mmol, 1.2 equiv.) was added dropwise at room temperature and the solution was stirred for 2 hours. The reaction was concentrated under reduced pressure and the residue was purified via flash-column chromatography (20% EtOAc/DCM) to afford the free alkyne as an off-white solid (194.9 mg, 54% yield).

This reaction did not use glassware that was flame- or oven-dried. The free alkyne (38.0 mg, 0.079 mmol, 1 equiv.) and LiOH·H_2_O (13.2 mg, 0.315 mmol, 4 equiv.) were added to a 2-dram vial, followed by THF (0.95 mL), H_2_O (0.32 mL), and EtOH (0.32 mL). The mixture was stirred at room temperature for 16 hours. The solution was then neutralized with 1 M HCl (pH paper) and the organic were concentrated *in vacuo*. The crude material was purified via flash column chromatography (1:1 PhMe:Acetone w/ 0.1% v/v AcOH, dry loaded with silica gel) to afford the alkynyl-quinazolinone free acid as a white solid (27.1 mg, 76% yield).

An oven-dried 2-dram vial was equipped with a magnetic stir bar and cooled to room temperature under a positive flow of argon gas. The alkynyl-quinazolinone free acid (11.3 mg, 0.025 mmol, 1 equiv.), CuI (1 mg, 0.005 mmol, 20 mol%), and TBTA (2.7 mg, 0.005 mmol, 20 mol%) were added sequentially to the vial. 0.13 mL of DMF followed by 0.13 mL of H_2_O was added to the solids, followed by the addition of biotin azide (250 μL of a 100 mM DMSO stock solution, 0.025 mmol, 1 equiv.). The reaction was placed in an oil bath pre-heated to 85 °C and stirred for 20 hours. The vial was removed from the oil bath and allowed to cool to room temperature. The crude material was filtered through a pad of celite eluting with EtOAc. The material was purified via reverse-phase column chromatography. MS (ESI+): m/z = 898.54 [M + H]+.

**HPLC traces of the compounds 9-32**

*3-(3-(2-(methylamino)-2-oxoethyl)-4-oxo-3,4-dihydroquinazolin-2-yl)propanoic acid,* ***9***

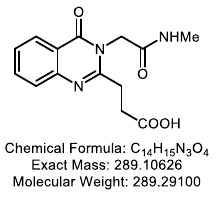

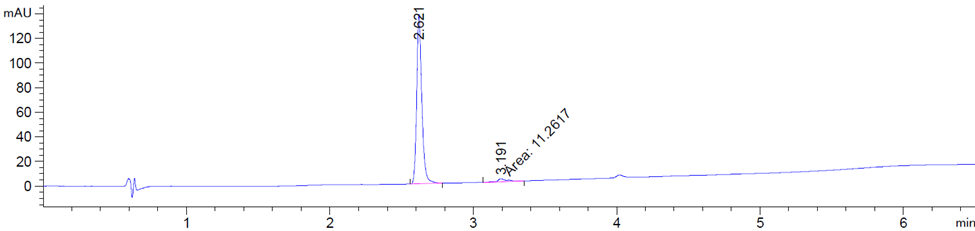

*3-(3-(2-(ethylamino)-2-oxoethyl)-4-oxo-3,4-dihydroquinazolin-2-yl)propanoic acid,* ***10***

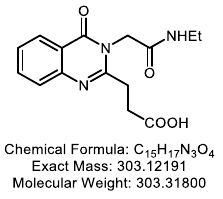

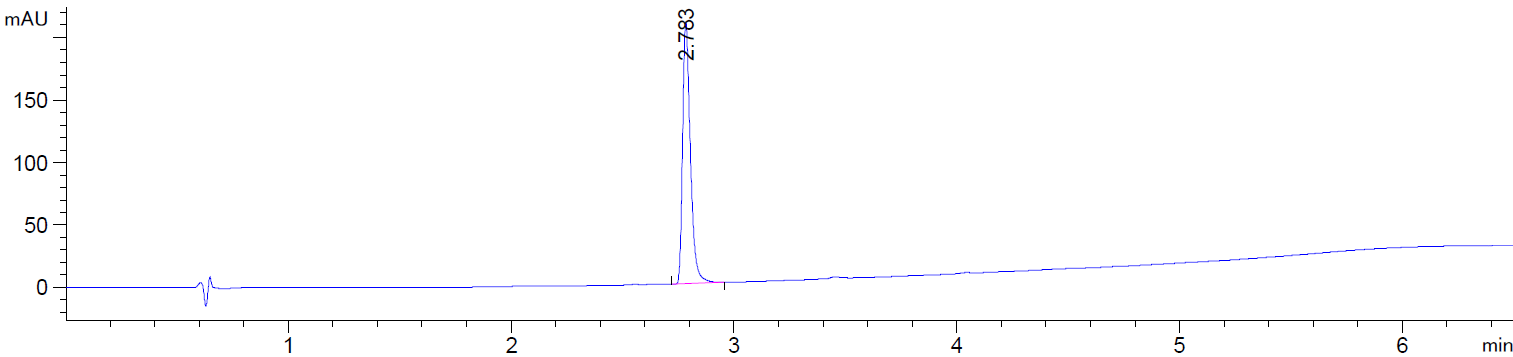

*3-(3-(2-(cyclopropylamino)-2-oxoethyl)-4-oxo-3,4-dihydroquinazolin-2-yl)propanoic acid,* ***11***

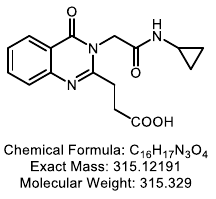

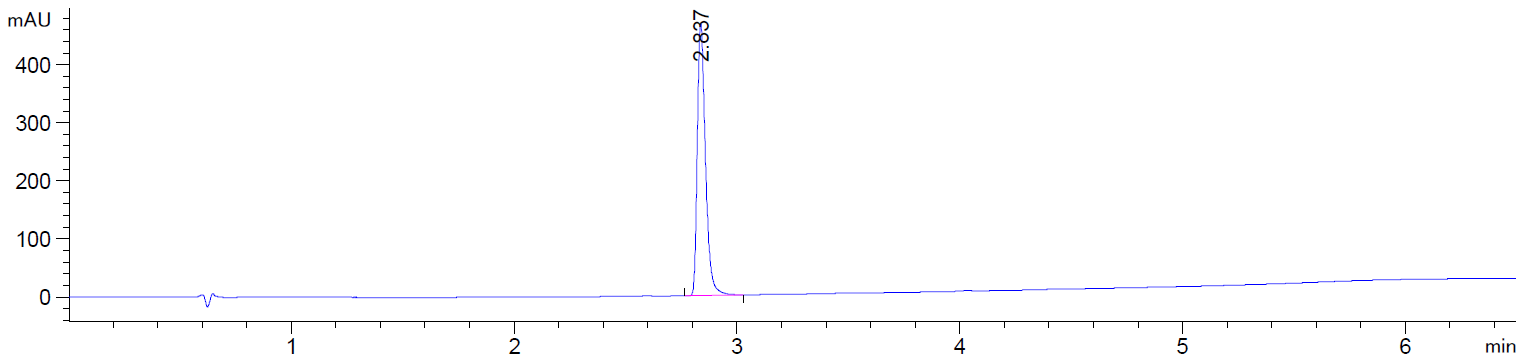

*3-(3-(2-(tert-butylamino)-2-oxoethyl)-4-oxo-3,4-dihydroquinazolin-2-yl)propanoic acid,* ***12***

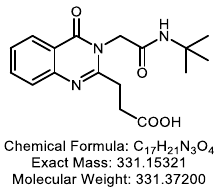

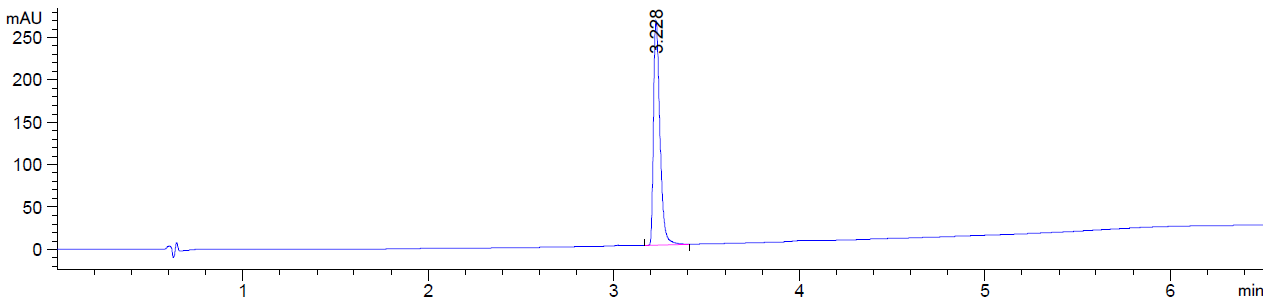

*3-(3-(2-(adamantan-1-yl)amino)-2-oxoethyl)-4-oxo-3,4-dihydroquinazolin-2-yl)propanoic acid,* ***13***

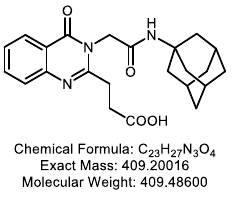

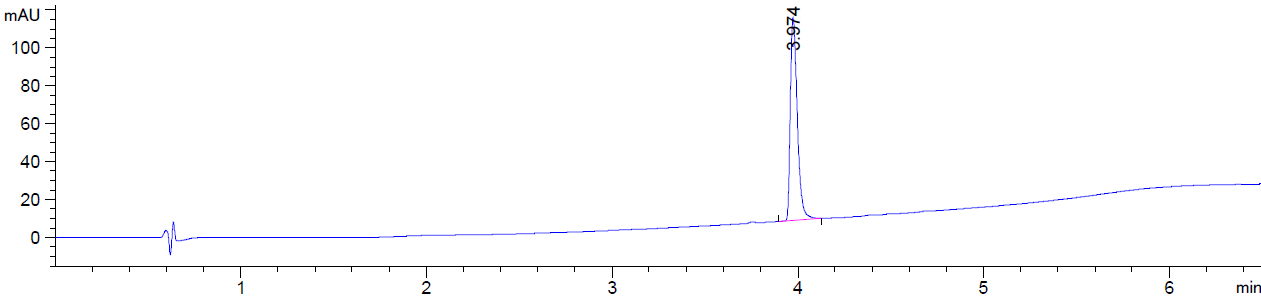

*3-(3-(2-((cyclohexylmethyl)amino)-2-oxoethyl)-4-oxo-3,4-dihydroquinazolin-2-yl)propanoic acid,* ***14***

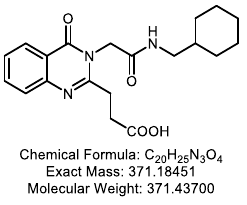

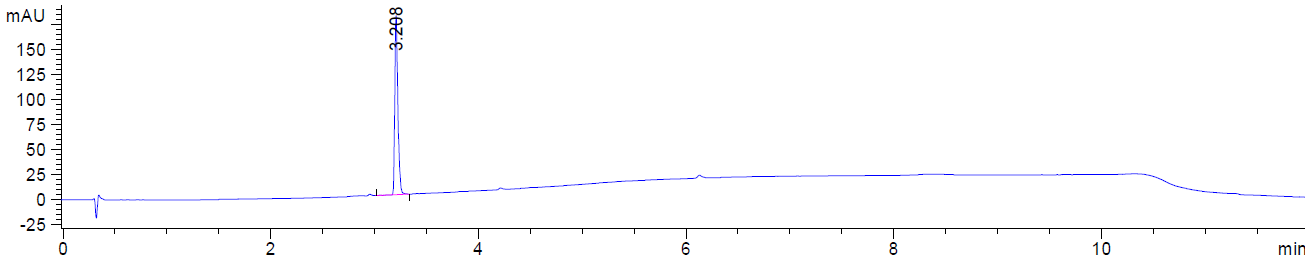

*3-(3-(2-(benzylamino)-2-oxoethyl)-4-oxo-3,4-dihydroquinazolin-2-yl)propanoic acid,* ***15***

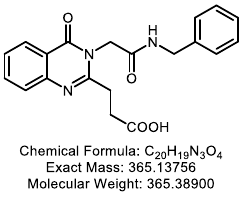

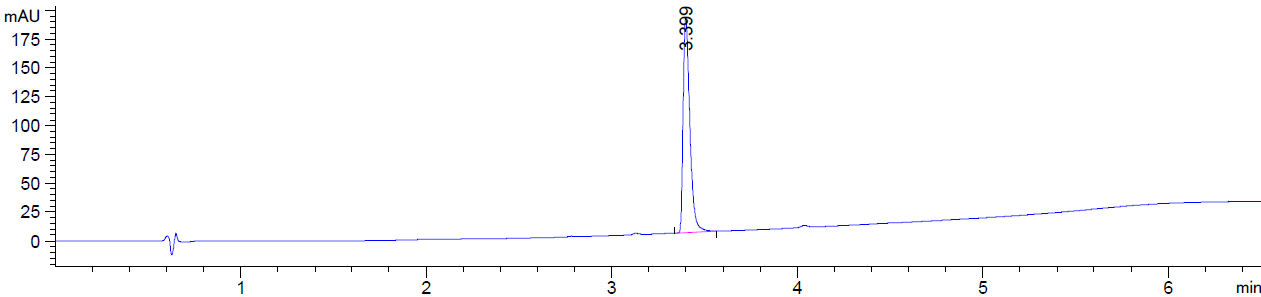

*3-(3-(2-((2-methoxybenzyl)amino)-2-oxoethyl)-4-oxo-3,4-dihydroquinazolin-2-yl)propanoic acid,* ***16***

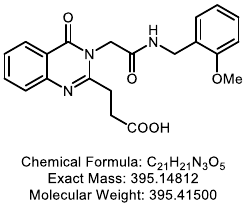

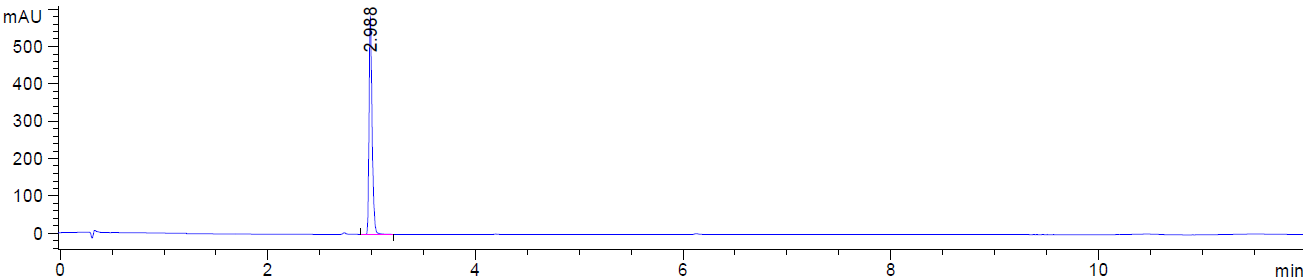

*3-(3-(2-((2-chlorobenzyl)amino)-2-oxoethyl)-4-oxo-3,4-dihydroquinazolin-2-yl)propanoic acid,* ***17***

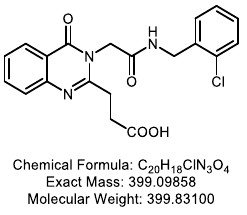

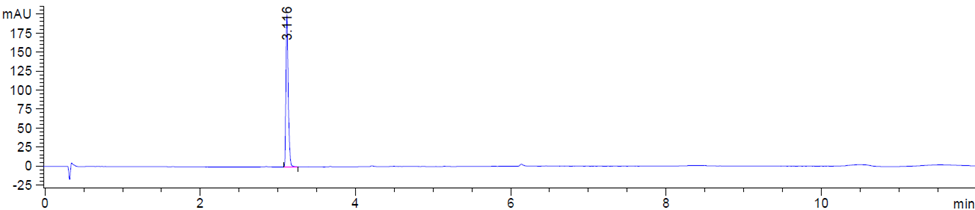

*3-(4-oxo-3-(2-oxo-2-((4-(trifluoromethyl)benzyl)amino)ethyl)-3,4-dihydroquinazolin-2-yl)propanoic acid,* ***18***

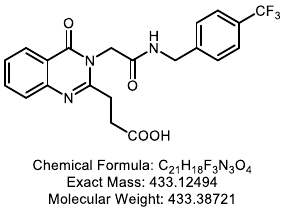

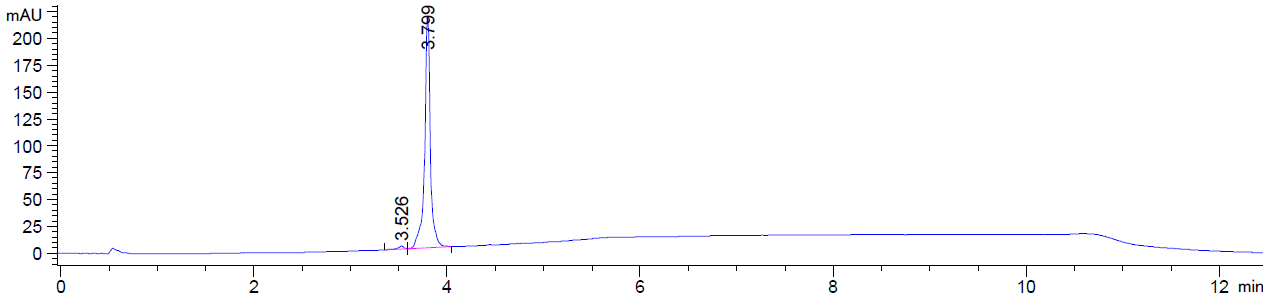

*3-(4-oxo-3-(2-oxo-2-((pyridin-3-ylmethyl)amino)ethyl)-3,4-dihydroquinazolin-2-yl)propanoic acid,* ***19***

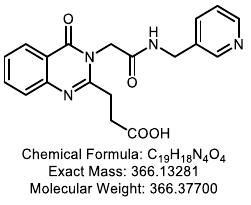

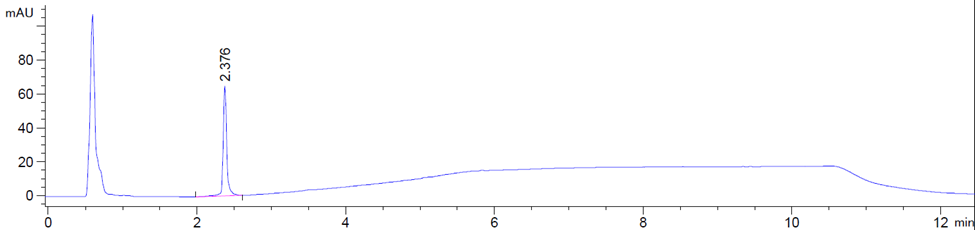

*3-(3-(2-((4-methoxybenzyl)amino)-2-oxoethyl)-4-oxo-3,4-dihydroquinazolin-2-yl)propanoic acid,* ***20***

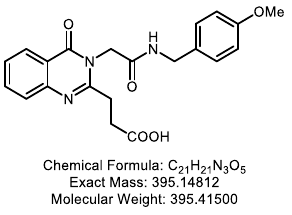

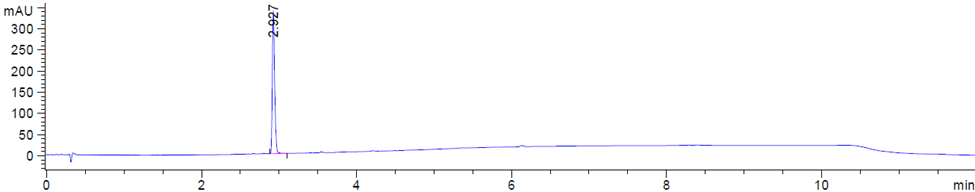

*3-(4-oxo-3-(2-oxo-2-(phenethylamino)ethyl)-3,4-dihydroquinazolin-2-yl)propanoic acid,* ***21***

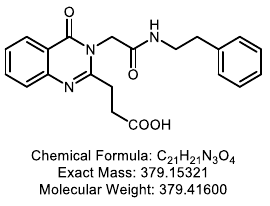

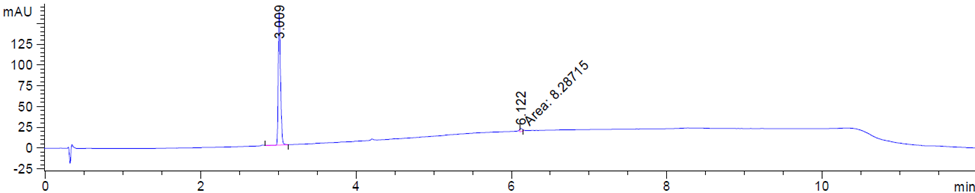

*3-(3-(2-((2-methoxyphenethyl)amino)-2-oxoethyl)-4-oxo-3,4-dihydroquinazolin-2-yl)propanoic acid,* ***22***

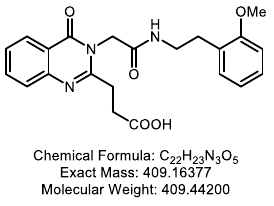

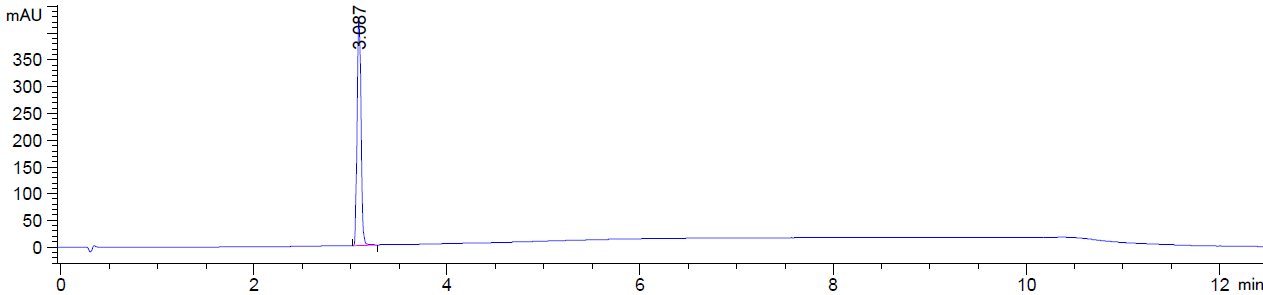

*3-(8-fluoro-3-(2-((2-methoxybenzyl)amino)-2-oxoethyl)-4-oxo-3,4-dihydroquinazolin-2-yl)propanoic acid,****23***

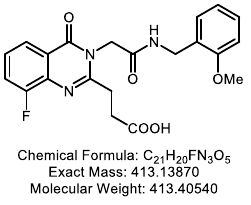

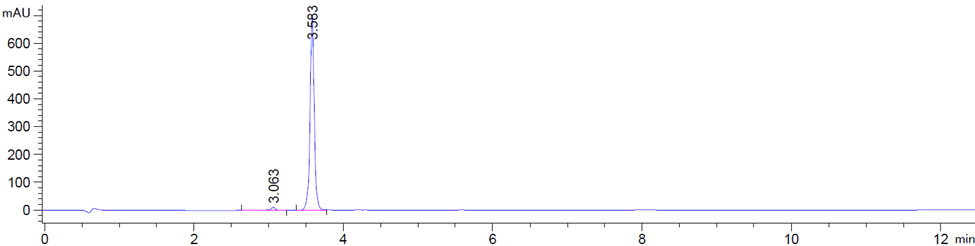

*3-(3-(2-((2-methoxybenzyl)amino)-2-oxoethyl)-8-methyl-4-oxo-3,4-dihydroquinazolin-2-yl)propanoic acid,* ***24***

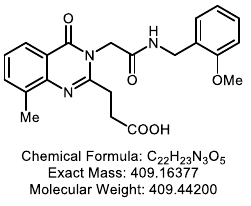

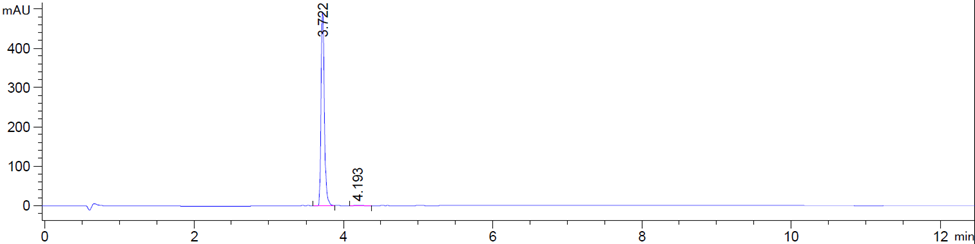

*3-(8-chloro-3-(2-((2-methoxybenzyl)amino)-2-oxoethyl)-4-oxo-3,4-dihydroquinazolin-2-yl)propanoic acid,* ***25***

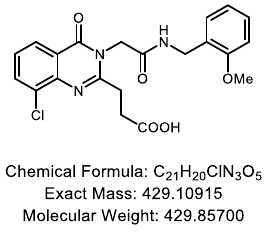

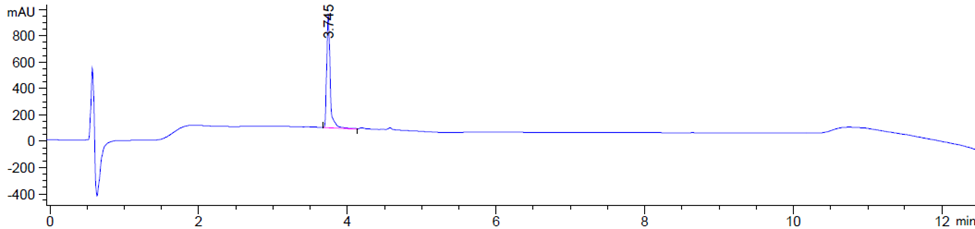

*3-(8-chloro-6-iodo-3-(2-((2-methoxybenzyl)amino)-2-oxoethyl)-4-oxo-3,4-dihydroquinazolin-2-yl)propanoic acid,* ***26***

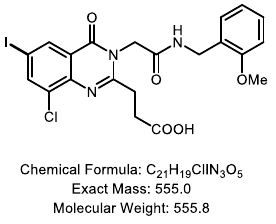

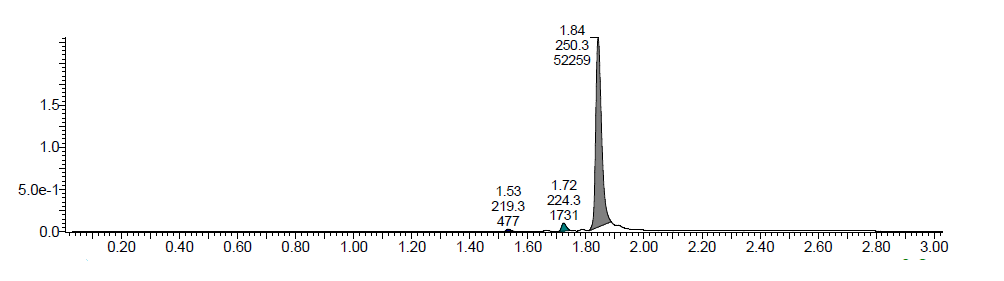

*3-(8-chloro-3-(2-((2-methylbenzyl)amino)-2-oxoethyl)-4-oxo-3,4-dihydroquinazolin-2-yl)propanoic acid,* ***27***

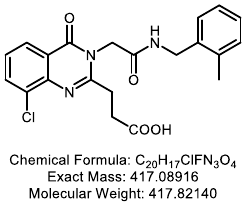

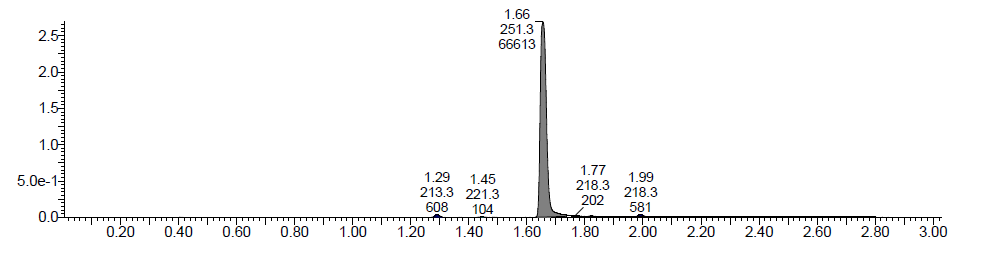

*3-(8-chloro-3-(2-((2-fluorobenzyl)amino)-2-oxoethyl)-4-oxo-3,4-dihydroquinazolin-2-yl)propanoic acid,* ***28***

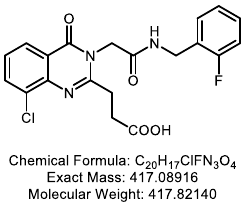

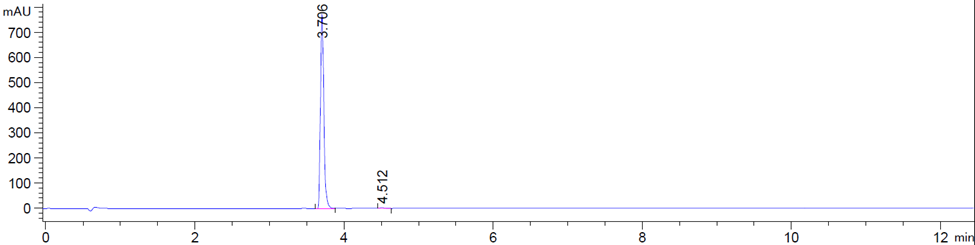

*3-(8-chloro-3-(2-((2-chlorobenzyl)amino)-2-oxoethyl)-4-oxo-3,4-dihydroquinazolin-2-yl)propanoic acid,* ***29***

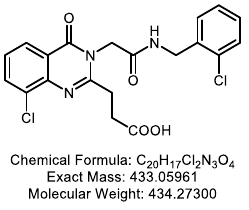

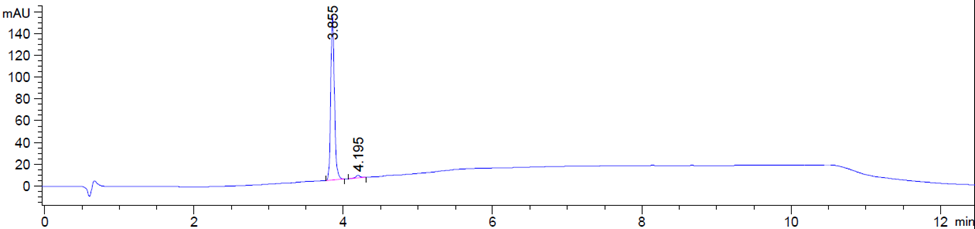

*3-(8-chloro-4-oxo-3-(2-oxo-2-(((tetrahydro-2H-pyran-2-yl)methyl)amino)ethyl)-3,4-dihydroquinazolin-2-yl)propanoic acid,* ***30***

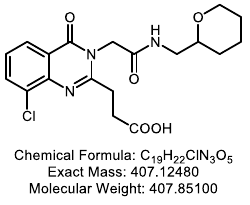

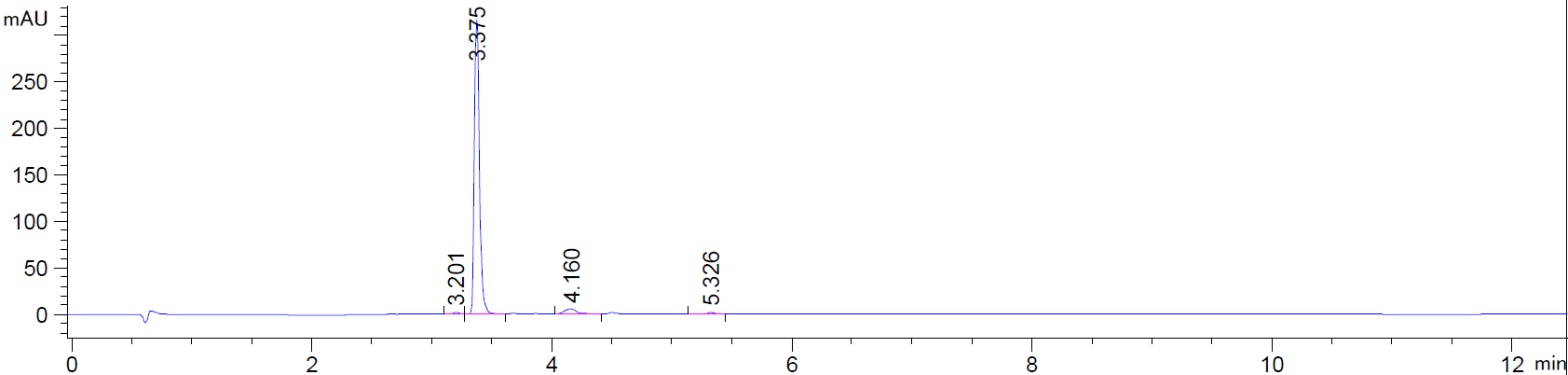

*3-(8-chloro-4-oxo-3-(2-oxo-2-(((tetrahydrofuran-2-yl)methyl)amino)ethyl)-3,4-dihydroquinazolin-2-yl)propanoic acid,* ***31***

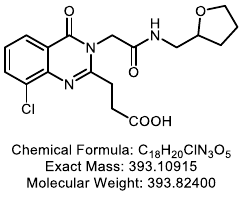

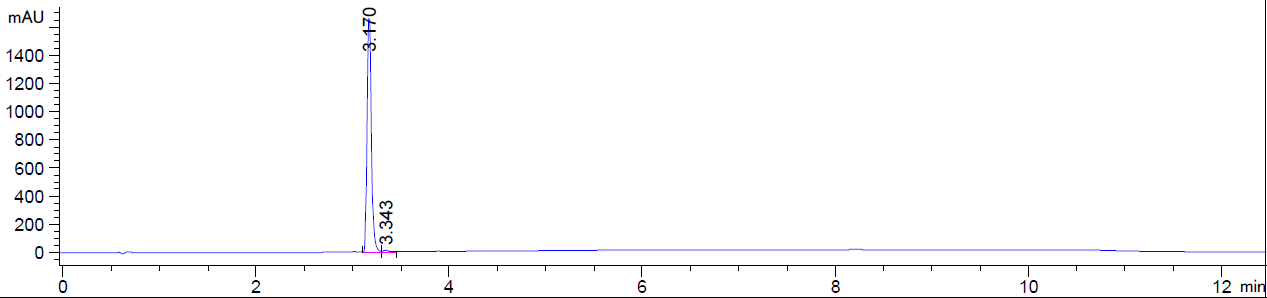

*3-(3-(2-(benzyl(methyl)amino)-2-oxoethyl)-4-oxo-3,4-dihydroquinazolin-2-yl)propanoic acid,* ***32***

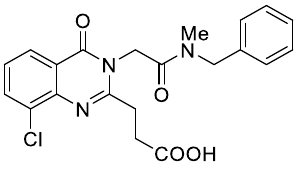

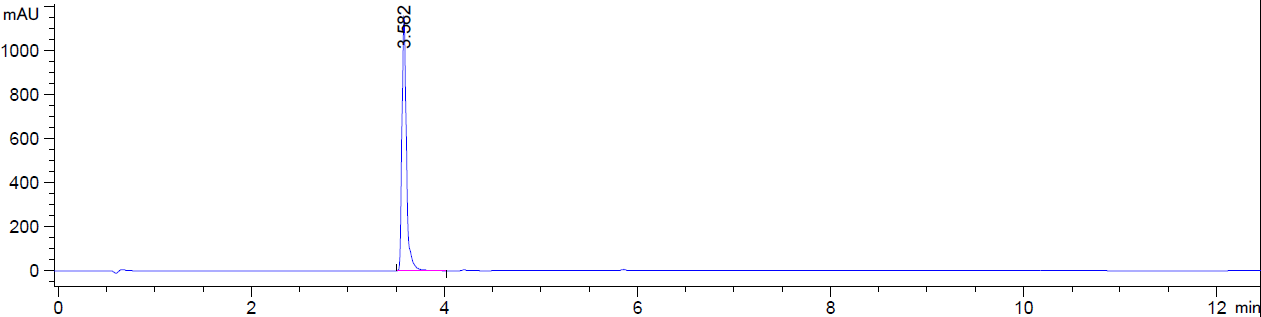

*3-(8-chloro-3-(2-((2-methoxybenzyl)amino)-2-oxoethyl)-4-oxo-6-(1-(13-oxo-17-((3aS,4S,6aR)-2-oxohexahydro-1H-thieno[3,4-d]imidazol-4-yl)-3,6,9-trioxa-12-azaheptadecyl)-1H-1,2,3-triazol-4-yl)-3,4-dihydroquinazolin-2-yl)propanoic acid,* ***33***

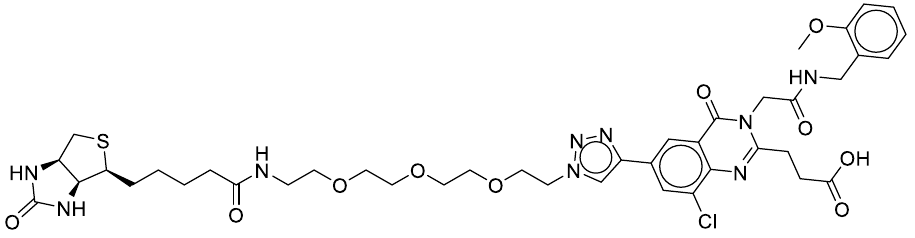

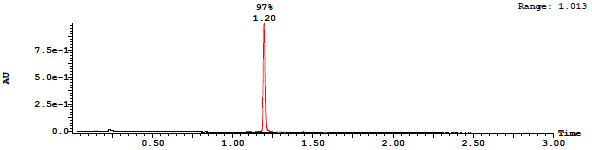
